# Anxiety and the neurobiology of temporally uncertain threat anticipation

**DOI:** 10.1101/2020.02.25.964734

**Authors:** Juyoen Hur, Jason F. Smith, Kathryn A. DeYoung, Allegra S. Anderson, Jinyi Kuang, Hyung Cho Kim, Rachael M. Tillman, Manuel Kuhn, Andrew S. Fox, Alexander J. Shackman

## Abstract

When extreme, anxiety—a state of distress and arousal prototypically evoked by uncertain danger—can be debilitating. Uncertain anticipation is a shared feature of situations that elicit signs and symptoms of anxiety across psychiatric disorders, species, and assays. Despite the profound significance of anxiety for human health and wellbeing, the neurobiology of uncertain-threat anticipation remains unsettled. Leveraging a paradigm adapted from animal research and optimized for functional MRI signal decomposition, we examined the neural circuits engaged during the anticipation of temporally uncertain and certain threat in 99 men and women. Results revealed that the neural systems recruited by uncertain and certain threat anticipation are anatomically co-localized in fronto-cortical regions, extended amygdala, and periaqueductal gray. Comparison of the threat conditions demonstrated that this circuitry can be fractionated, with fronto-cortical regions showing relatively stronger engagement during the anticipation of uncertain threat, and the extended amygdala showing the reverse pattern. Although there is widespread agreement that the bed nucleus of the stria terminalis and dorsal amygdala—the two major subdivisions of the extended amygdala—play a critical role in orchestrating adaptive responses to potential danger, their precise contributions to human anxiety have remained contentious. Follow-up analyses demonstrated that these regions show statistically indistinguishable responses to temporally uncertain and certain threat anticipation. These observations provide a framework for conceptualizing anxiety and fear, for understanding the functional neuroanatomy of threat anticipation in humans, and for accelerating the development of more effective intervention strategies for pathological anxiety.

**SIGNIFICANCE STATEMENT:** Anxiety—an emotion prototypically associated with the anticipation of uncertain harm—has profound significance for public health, yet the underlying neurobiology remains unclear. Leveraging a novel neuroimaging paradigm in a relatively large sample, we identify a core circuit responsive to both uncertain and certain threat anticipation, and show that this circuitry can be fractionated into subdivisions with a bias for one kind of threat or the other. The extended-amygdala occupies center-stage in neuropsychiatric models of anxiety, but its functional architecture has remained contentious. Here we demonstrate that its major subdivisions show statistically indistinguishable responses to temporally uncertain and certain threat. Collectively, these observations indicate the need to revise how we think about the neurobiology of anxiety and fear.

**RESOURCE SHARING:** Raw data are available at the National Institute of Mental Health’s Data Archive. Key statistical maps are or will be publicly available at NeuroVault.org.

## INTRODUCTION

Anxiety is widely conceptualized as a state of distress and arousal elicited by the anticipation of uncertain danger (Davis et al., 2010; Grupe and Nitschke, 2013). Anxiety lies on a continuum and, when extreme, can be debilitating (Salomon et al., 2015; Conway et al., 2019). Anxiety disorders are the most common family of psychiatric illnesses and existing treatments are inconsistently effective or associated with significant adverse effects, underscoring the urgency of developing a clearer understanding of the underlying neurobiology (Griebel and Holmes, 2013; Global Burden of Disease Collaborators, 2016; Craske et al., 2017).

Perturbation and recording studies in mice have begun to reveal the specific molecules and cellular ensembles that underlie defensive responses to uncertain threat (Fadok et al., 2018; Fox and Shackman, 2019), but the relevance of these tantalizing discoveries to the complexities of human anxiety is unclear. Humans and mice diverged ∼75 MYA, leading to marked behavioral, genetic, and neurobiological differences between the two species (Van Essen et al., 2019). The role of fronto-cortical regions that are especially well-developed in humans—including the midcingulate cortex (MCC), anterior insula (AI), and dorsolateral prefrontal cortex (dlPFC)—also remains opaque, reflecting equivocal or absent anatomical homologies and the use of disparate paradigms across species (Vogt and Paxinos, 2014; Shackman et al., 2016; Carlén, 2017; Roberts, 2020).

Beneath the neocortex, the role of the central extended amygdala—including the dorsal amygdala in the region of the central nucleus (Ce) and the bed nucleus of the stria terminalis (BST)—remains particularly contentious (Fox and Shackman, 2019). Inspired by an earlier-generation of lesion studies in rodents (Davis, 2006), it is widely believed that these regions are functionally dissociable, with the amygdala mediating phasic responses to clear-and-immediate danger (‘acute threat’) and the BST mediating sustained responses to uncertain-or-remote danger (‘potential threat’) (e.g., Sylvers et al., 2011; Somerville et al., 2013; Avery et al., 2016; LeDoux and Pine, 2016; Klumpers et al., 2017; Watson et al., 2017). This ‘strict-segregation’ hypothesis has even been enshrined in the National Institute of Mental Health’s (NIMH) Research Domain Criteria (RDoC) framework (National Institute of Mental Health, 2011, 2020a, b). Yet, a growing body of optogenetic, chemogenetic, and electrophysiological work in rodents demonstrates that defensive responses elicited by the anticipation of uncertain threat (e.g. elevated-plus maze) are assembled by microcircuits encompassing both regions (Gungor and Paré, 2016; Lange et al., 2017; Ahrens et al., 2018; Pomrenze et al., 2019a; Pomrenze et al., 2019b; Ressler et al., 2020; Griessner et al., *in press*), motivating the competing hypothesis that the dorsal amygdala and BST are both important substrates for human anxiety (Shackman and Fox, 2016; Fox and Shackman, 2019).

To address these fundamental questions, we combined fMRI with a novel threat-anticipation task in 99 adults. Advanced data acquisition and processing techniques enhanced resolution of subcortical regions. Building on earlier work (e.g., Somerville et al., 2013; Grupe et al., 2016), the Maryland Threat Countdown (MTC) paradigm is an fMRI-optimized variant of assays that have been validated using fear-potentiated startle and acute pharmacological manipulations in rodents (Miles et al., 2011; Daldrup et al., 2015), and humans (Hefner et al., 2013), maximizing translational relevance. It takes the form of a 2 (*Valence:* Threat/Safety) × 2 (*Temporal Certainty:* Uncertain/Certain) randomized event-related design (**Fig. 1**). On Certain Threat trials, subjects saw a descending stream of integers for 18.75 s, sufficiently long to enable the dissection of onset-evoked from sustained hemodynamic responses. To ensure robust emotion, this anticipatory epoch (‘countdown’) always culminated with the delivery of a multi-modal reinforcer (aversive shock, photograph, and audio-clip). Uncertain Threat trials were similar, but the integer stream was randomized and presented for an uncertain and variable duration (*M*=18.75 s, Range=8.75-30.00). Here, subjects knew the threat was coming, but they did not know *when* it would occur. Safety trials were similar, but terminated in benign reinforcers. Comparison of the well-matched anticipatory epochs enabled us to rigorously isolate circuits recruited during uncertain-threat anticipation.

**Figure 1.**
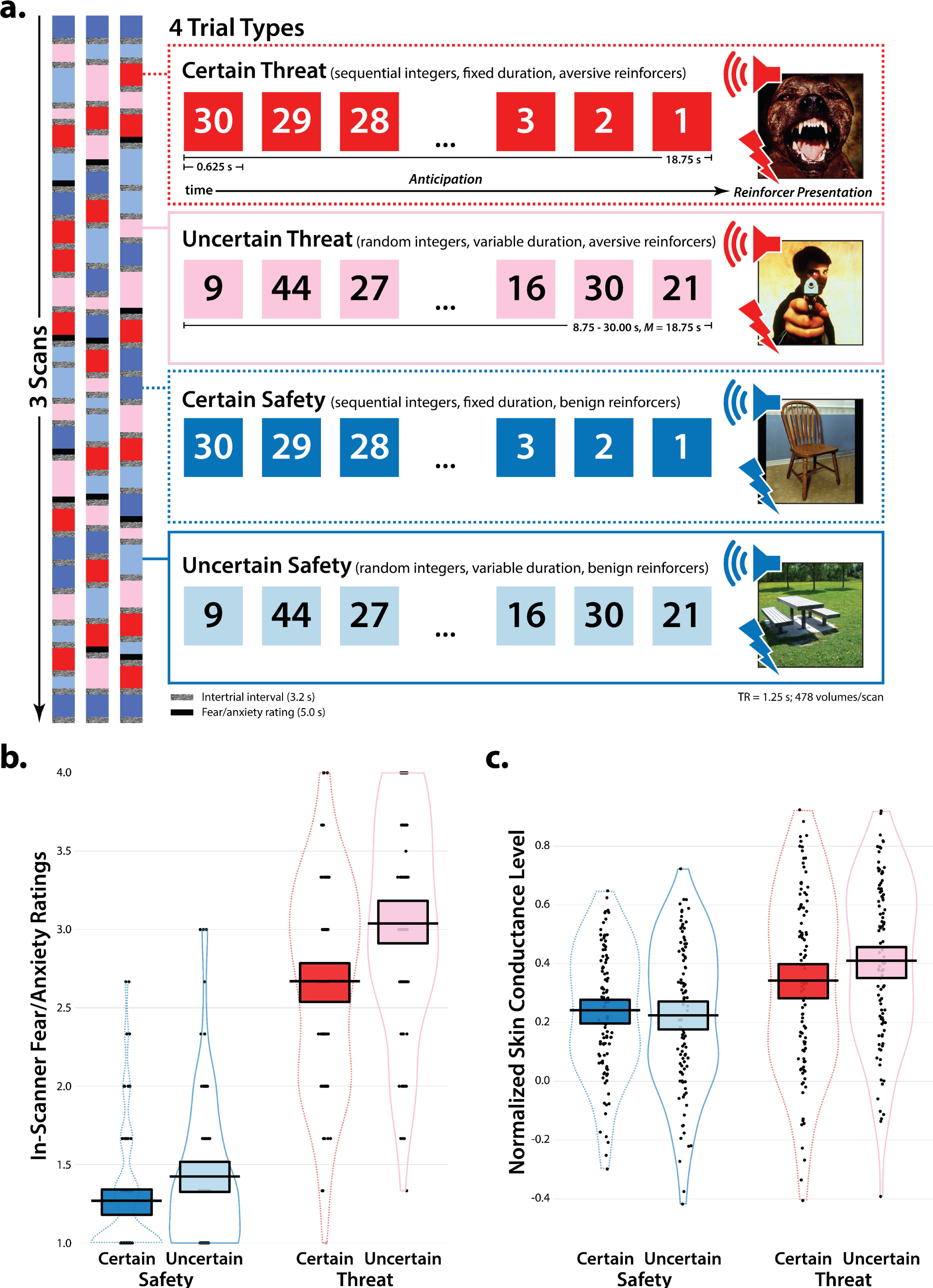
Maryland Threat Countdown (MTC) Paradigm. As shown schematically in panel ***a***, the MTC paradigm takes the form of a 2 (*Valence:* Threat/Safety) × 2 (*Temporal Certainty:* Uncertain/Certain) repeated-measures design. See the main text for a general description and Materials and Methods for details. Subjects provided ratings of anticipatory fear/anxiety for each trial type during each scan. Skin conductance was continuously acquired during scanning. Simulations were used to optimize the detection and deconvolution of task-related hemodynamic signals (variance inflation factors <1.54). Central panels depict the structure of each trial type. Trial valence was continuously signaled during the anticipatory epoch by the background color of the display. Safety trials were similar, but terminated with the delivery of benign stimuli (e.g. just-perceptible electrical stimulation). Trial certainty was signaled by the nature of the integer stream. Certain trials always began with the presentation of 30. On Uncertain trials, integers were randomly drawn from a uniform distribution ranging from 1 to 45 to reinforce the belief that uncertain trials could be much longer than certain ones. To mitigate potential confusion and eliminate mnemonic demands, a lower-case ‘c’ or ‘u’ was presented at the lower edge of the display throughout the anticipatory epoch (not depicted). As shown in panels ***b*** and ***c***, threat anticipation robustly increased subjective symptoms (in-scanner ratings) and objective signs (skin conductance) of anxiety, and this was particularly evident when the timing of aversive stimulation was uncertain (Valence × Certainty, *p*s<.001; Uncertain Threat > Certain Threat, *p*s<.001). Panels *b* and *c* depict the data (*black points; individual participants*), density distribution (*bean plots*), Bayesian 95% highest density interval (HDI; *colored bands*), and mean (*black bars*) for each condition. HDIs permit population-generalizable visual inferences about mean differences and were estimated using 1,000 samples from a posterior Gaussian distribution. Abbreviations—TR, repetition time (i.e. the time required to collect a single volume of fMRI data).

## MATERIALS AND METHODS

### Subjects

As part of an on-going prospective-longitudinal study focused on the emergence of anxiety disorders and depression, we used well-established measures of dispositional negativity (often termed neuroticism or negative emotionality; Shackman et al., 2018; Hur et al., 2019; Hur et al., *in press*) to screen 6,594 young adults (57.1% female; 59.0% White, 19.0% Asian, 9.9% African American, 6.3% Hispanic, 5.8% Multiracial/Other; *M*=19.2 years, *SD*=1.1 years). Screening data were stratified into quartiles (top quartile, middle quartiles, bottom quartile) separately for men and women. Individuals who met preliminary inclusion criteria were independently recruited from each of the resulting six strata. Given the focus of the larger study, approximately half the subjects were recruited from the top quartile, with the remainder split between the middle and bottom quartiles (i.e., 50% high, 25% medium, and 25% low), enabling us to sample a wide range of risk for the development of internalizing disorders. A total of 121 subjects were recruited. Of these, 2 withdrew during the imaging assessment due to excess distress. Of the 119 subjects who completed the imaging assessment, a total of 20 were excluded from analyses due to incidental neurological findings (*n*=4), scanner problems (*n*=2), insufficient fMRI data (<2 usable scans, *n*=1), excessive global motion artifact (see below; *n*=3), or excessive task-correlated motion (see below, *n*=10). This yielded a final sample of 99 subjects (52 females; 65.7% White, 17.2% Asian, 8.1% African American, 3.0% Hispanic, 6.1% Multiracial/Other; *M*=18.8 years, *SD*=0.4 years), providing substantially greater power to detect medium-sized (0.5 < Cohen’s *d* < 0.8) statistical effects (Geuter et al., 2018) compared to typical fMRI studies of uncertain threat anticipation (median N=29; range=15-108; Chavanne and Robinson, in press). All subjects had normal or corrected-to-normal color vision; and reported the absence of lifetime neurological symptoms, pervasive developmental disorder, very premature birth, medical conditions that would contraindicate MRI, and prior experience with noxious electrical stimulation. All subjects were free from a lifetime history of psychotic and bipolar disorders; a current diagnosis of a mood, anxiety, or trauma disorder (past 2 months); severe substance abuse; active suicidality; and on-going psychiatric treatment as determined by an experienced masters-level diagnostician using the Structured Clinical Interview for DSM-5 (First et al., 2015). Subjects provided informed written consent and all procedures were approved by the Institutional Review Board at the University of Maryland, College Park.

### Maryland Threat Countdown (MTC) fMRI Paradigm

#### Paradigm Structure and Design Considerations

Building on earlier imaging work (Somerville et al., 2013; Grupe et al., 2016; Pedersen et al., 2019), the Maryland Threat Countdown (MTC) paradigm is an fMRI-optimized version of temporally uncertain-threat assays that have been validated using fear-potentiated startle and acute anxiolytic administration (e.g. benzodiazepine) in mice (Daldrup et al., 2015; Lange et al., 2017), rats (Miles et al., 2011), and humans (Hefner et al., 2013), enhancing its translational relevance. The MTC paradigm takes the form of a 2 (*Valence:* Threat/Safety) × 2 (*Temporal Certainty:* Uncertain/Certain) randomized event-related design (3 scans; 6 trials/condition/scan). Simulations were used to optimize the detection and deconvolution of task-related hemodynamic signals (variance inflation factors <1.54). Stimulus presentation and ratings acquisition were controlled using Presentation software (version 19.0, Neurobehavioral Systems, Berkeley, CA).

On Certain Threat trials, subjects saw a descending stream of integers (‘count-down;’ e.g. 30, 29, 28…3, 2, 1) for 18.75 s. To ensure robust emotion, this anticipatory epoch always culminated with the delivery of a noxious electric shock, unpleasant photographic image (e.g. mutilated body), and thematically related audio clip (e.g. scream, gunshot). Uncertain Threat trials were similar, but the integer stream was randomized and presented for an uncertain and variable duration (8.75-30.00 s; *M*=18.75 s). Here, subjects knew that something aversive was going to occur, but they had no way of knowing precisely when it would occur. Consistent with recent recommendations (Shackman and Fox, 2016), the average duration of the anticipatory epoch was identical across conditions, ensuring an equal number of measurements (TRs/condition). Mean duration was chosen to enhance detection of task-related differences in the blood oxygen level-dependent (BOLD) signal (Henson, 2007), and to enable dissection of onset from genuinely sustained responses. Safety trials were similar, but terminated with the delivery of benign reinforcers (see below). Valence was continuously signaled during the anticipatory epoch by the background color of the display. Temporal certainty was signaled by the nature of the integer stream. Certain trials always began with the presentation of the number 30 (**Fig. 1**). On uncertain trials integers were randomly drawn from a near-uniform distribution ranging from 1 to 45 to reinforce the impression that Uncertain trials could be much longer than Certain ones and to minimize incidental temporal learning (‘time-keeping’). To mitigate potential confusion and eliminate mnemonic demands, a lower-case ‘c’ or ‘u’ was presented at the lower edge of the display throughout the anticipatory epoch. White-noise visual masks (3.2 s) were presented between trials to minimize persistence of the visual reinforcers in iconic memory. Subjects provided ratings of anticipatory fear/anxiety for each trial type during each scan using an MRI-compatible response pad (MRA, Washington, PA; **Fig. 1**). Subjects were instructed to rate the intensity of the fear/anxiety experienced during the prior anticipatory (‘countdown’) epoch using a 1 (*minimal*) to 4 (*maximal*) scale. Subjects were promoted to rate each trial type once per scan. A total of 6 additional echo-planar imaging (EPI) volumes were acquired at the beginning and end of each scan (see below).

#### Procedures

Prior to scanning, subjects practiced an abbreviated version of the paradigm—without electrical stimulation—until they indicated and staff confirmed that they understood the task. Benign and aversive electrical stimulation levels were individually titrated.

#### Benign Stimulation

Subjects were asked whether they could “reliably detect” a 20 V stimulus and whether it was “at all unpleasant.” If the subject could not detect the stimulus, the voltage was increased by 4 V and the process repeated. If the subject indicated that the stimulus was unpleasant, the voltage was reduced by 4V and the process repeated. The final level chosen served as the benign electrical stimulation during the imaging assessment (*M*=20.67, *SD*=6.23).

#### Aversive Stimulation

Subjects received a 100 V stimulus and were asked whether it was “as unpleasant as you are willing to tolerate.” If the subject indicated that they were willing to tolerate more intense stimulation, the voltage was increased by 10 V and the process repeated. If the subject indicated that the stimulus was too intense, the voltage was reduced by 5 V and the process repeated. The final level chosen served as the aversive electrical stimulation during the imaging assessment (*M*=115.21, *SD*=25.05). Following each scan of the MTC paradigm, we re-assessed whether stimulation was sufficiently intense and re-calibrated as necessary. In total, 32.3% of subjects adjusted the level of benign or aversive stimulation at least once during the imaging assessment.

#### Electrical Stimuli

Electrical stimuli (100 ms; 2 ms pulses every 10 ms) were generated using an MRI-compatible constant-voltage stimulator system (STMEPM-MRI; Biopac Systems, Inc., Goleta, CA). Stimuli were delivered using MRI-compatible, disposable carbon electrodes (Biopac) attached to the fourth and fifth phalanges of the non-dominant hand.

#### Visual Stimuli

Visual stimuli (1.8 s) were digitally back-projected (Powerlite Pro G5550, Epson America, Inc., Long Beach, CA) onto a semi-opaque screen mounted at the head-end of the scanner bore and viewed using a mirror mounted on the head-coil. A total of 72 photographs were selected from the International Affective Picture System (IAPS identification numbers)—*Benign:* 1670, 2026, 2038, 2102, 2190, 2381, 2393, 2397, 2411, 2850, 2870, 2890, 5390, 5471, 5510, 5740, 7000, 7003, 7004, 7014, 7020, 7026, 7032, 7035, 7050, 7059, 7080, 7090, 7100, 7140, 7187, 7217, 7233, 7235, 7300, 7950. *Aversive:* 1300, 3000, 3001, 3010, 3015, 3030, 3051, 3053, 3061, 3062, 3063, 3069, 3100, 3102, 3150, 3168, 3170, 3213, 3400, 3500, 6022, 6250, 6312, 6540, 8230, 9042, 9140, 9253, 9300, 9405, 9410, 9414, 9490, 9570, 9584, 9590. (Lang et al., 2008). Based on normative ratings, the aversive images were significantly more negative and arousing than the benign images, *t*(70)>24.3, *p*<.001. On a 1 (*negative/low-arousal*) to 9 (*positive/high-arousal*) scale, the mean valence and arousal scores were 2.2 (*SD*=0.6) and 6.3 (*SD*=0.6) for the aversive images, and 5.2 (*SD*=0.4) and 2.8 (*SD*= 0.3) for the benign images.

#### Auditory Stimuli

Auditory stimuli (0.80 s) were delivered using an amplifier (PA-1 Whirlwind) with in-line noise-reducing filters and ear buds (S14; Sensimetrics, Gloucester, MA) fitted with noise-reducing ear plugs (Hearing Components, Inc., St. Paul, MN). A total of 72 auditory stimuli (half aversive, half benign) were adapted from open-access online sources.

### Peripheral Physiology Data Acquisition

Peripheral physiology was continuously acquired during each fMRI scan using a Biopac system (MP-150). Skin conductance (250 Hz; 0.05 Hz high-pass) was measured using MRI-compatible disposable electrodes (EL507) attached to the second and third phalanges of the non-dominant hand. For imaging analyses, measures of respiration and breathing were also acquired using a respiration belt and photo-plethysmograph (first phalange of the non-dominant hand).

### MRI Data Acquisition

MRI data were acquired using a Siemens Magnetom TIM Trio 3 Tesla scanner (32-channel head-coil). Foam inserts were used to immobilize the participant’s head within the head-coil and mitigate potential motion artifact. Subjects were continuously monitored from the control room using an MRI-compatible eye-tracker (Eyelink 1000; SR Research, Ottawa, Ontario, Canada). Head motion was monitored using the AFNI real-time plugin (Cox, 1996). Sagittal T1-weighted anatomical images were acquired using a magnetization prepared rapid acquisition gradient echo (MPRAGE) sequence (TR=2,400 ms; TE=2.01 ms; inversion time=1060 ms; flip angle=8°; sagittal slice thickness=0.8 mm; in-plane=0.8 × 0.8 mm; matrix=300 × 320; field-of-view=240 × 256). A T2-weighted image was collected co-planar to the T1-weighted image (TR=3,200 ms; TE=564 ms; flip angle=120°). To enhance resolution, a multi-band sequence was used to collect oblique-axial echo planar imaging (EPI) volumes (multiband acceleration=6; TR=1,250 ms; TE=39.4 ms; flip angle=36.4°; slice thickness=2.2 mm, number of slices=60; in-plane resolution=2.1875 × 2.1875 mm; matrix=96 × 96). Images were collected in the oblique axial plane (approximately −20° relative to the AC-PC plane) to minimize potential susceptibility artifacts. Three 478-volume EPI scans were acquired. The scanner automatically discarded 7 volumes prior to the first recorded volume. To enable fieldmap correction, two oblique-axial spin echo (SE) images were collected in each of two opposing phase-encoding directions (rostral-to-caudal and caudal-to-rostral) at the same location and resolution as the functional volumes (i.e., co-planar; TR=7,220 ms; TE=73 ms). Following the last scan, subjects were removed from the scanner, debriefed, compensated, and discharged.

### Skin Conductance Data Pipeline

Skin conductance data were processed using PsPM (version 4.0.2) and in-house Matlab code (Bach and Friston, 2013; Bach et al., 2018). Data from each scan were band-pass filtered (0.01-0.25 Hz), resampled to match the TR used for fMRI data acquisition (1.25 s), and *z*-transformed. Using standard Matlab functions, SCR data were modeled in a manner that approximated that used for the fMRI data. A GLM was used to estimate skin conductance levels during the anticipatory epoch of each condition of the MTC paradigm (see above) for each subject (Bach et al., 2009; Bach et al., 2013; Bach, 2014). Predictors from the first-level fMRI model (see below) were convolved with a canonical skin conductance response function (Bach et al., 2010; Gerster et al., 2018), bandpass filtered to match the data, and *z*-transformed.

### MRI Data Pipeline

Methods were optimized to minimize spatial normalization error and other potential sources of noise. Structural and functional MRI data were visually inspected before and after processing for quality assurance.

#### Anatomical Data Processing

Methods were similar to those described in other recent reports by our group (Hur et al., 2018; Smith et al., 2018; Tillman et al., 2018). T1-weighted images were inhomogeneity corrected using N4 (Tustison et al., 2010) and filtered using the denoise function in ANTS (Avants et al., 2011). The brain was then extracted using a variant of the BEaST algorithm (Eskildsen et al., 2012) with brain-extracted and normalized reference brains from the IXI database (https://brain-development.org/ixi-dataset). Brain-extracted T1 images were normalized to a version of the brain-extracted 1-mm T1-weighted MNI152 (version 6) template (Grabner et al., 2006) modified to remove extracerebral tissue. This was motivated by evidence that brain-extracted T1 images and a brain-extracted template enhance the quality of spatial normalization (Fein et al., 2006; Acosta-Cabronero et al., 2008; Fischmeister et al., 2013). Normalization was performed using the diffeomorphic approach implemented in SyN (version 1.9.x.2017-09.11; Klein et al., 2009; Avants et al., 2011). T2-weighted images were rigidly co-registered with the corresponding T1 prior to normalization and the brain extraction mask from the T1 was applied. Tissue priors (Lorio et al., 2016) were unwarped to the native space of each T1 using the inverse of the diffeomorphic transformation. Brain-extracted T1 and T2 images were simultaneously segmented using native-space priors generated using FAST (FSL version 5.0.9) (Zhang et al., 2001) for use in T1-EPI co-registration (see below).

#### Fieldmap Data Processing

SE images were used to create a fieldmap in topup (Andersson et al., 2003; Smith et al., 2004; Graham et al., 2017). Fieldmaps were converted to radians, median filtered, and smoothed (2-mm). The average of the distortion-corrected SE images was inhomogeneity-corrected using N4, and brain-masked using 3dSkullStrip in AFNI (version 17.2.10; Cox, 1996). The resulting mask was minimally eroded to further exclude extracerebral voxels.

#### Functional Data Processing

EPI files were de-spiked using 3dDespike and slice-time corrected (to the center of the TR) using 3dTshift, inhomogeneity corrected using N4, and motion corrected to the first volume using a 12-parameter affine transformation implemented in ANTs. Recent work indicates that de-spiking is more effective than ‘scrubbing’ for attenuating motion-related artifacts (Jo et al., 2013; Siegel et al., 2014; Power et al., 2015). Transformations were saved in ITK-compatible format for subsequent use. The first volume was extracted for EPI-T1 co-registration. The reference EPI volume was simultaneously co-registered with the corresponding T1-weighted image in native space and corrected for geometric distortions using boundary-based registration (Greve and Fischl, 2009). This step incorporated the previously created fieldmap, undistorted SE, T1, white matter (WM) image, and masks. The spatial transformations necessary to transform each EPI volume from native space to the reference EPI, from the reference EPI to the T1, and from the T1 to the template were concatenated and applied to the processed (de-spiked and slice-time corrected) EPI data in a single step to minimize incidental spatial blurring. Normalized EPI data were resampled to 2-mm isotopic voxels using fifth-order b-splines and smoothed (6-mm FWHM) using 3DblurInMask.

#### Data Exclusions

To assess residual motion artifact, we computed the number of times the brain showed a volume-to-volume displacement >0.5 mm using the motion-corrected data. Scans with excess artifact (≥7.5%) were discarded. Three subjects with insufficient usable data (<2 scans) were excluded from analyses, while 6 subjects with 2 usable scans were retained. To assess task-correlated motion, we computed correlations between the design matrix and the motion estimates (see above). Scans showing extreme correlations (>2 *SD*) were discarded. On this basis, ten subjects with insufficient usable data (<2 scans) were excluded from analyses, while 19 subjects with 2 usable scans were retained.

#### Canonical First-Level fMRI Modeling

Modeling was performed using SPM12 (version 6678; https://www.fil.ion.ucl.ac.uk/spm). Band-pass was set to the hemodynamic response function (HRF) and 128 s for low and high pass, respectively. The MTC paradigm was modeled using variable-duration rectangular (‘box-car’) regressors time-locked to the anticipatory epochs of the Uncertain Threat, Certain Threat, and Uncertain Safety trials. Certain Safety trials were treated as an unmodeled (‘implicit’) high-level baseline. EPI volumes collected before the first trial, during intertrial intervals, and following the final trial were also unmodeled, and contributed to the baseline estimate. Regressors were convolved with a canonical HRF and its temporal derivative. The periods corresponding to the delivery of the four reinforcers and rating trials were modeled using a similar approach (**Fig. 1**). Volume-to-volume displacement and motion parameters (including 1- and 2-volume lagged versions) were also included, similar to other recent work (Reddan et al., 2018). To further attenuate potential noise, cerebrospinal fluid (CSF) time-series, instantaneous pulse and respiration rates, and their estimated effect on the BOLD time-series were also included as nuisance variates. ICA-AROMA (Pruim et al., 2015) was used to model several other potential sources of noise (brain-edge, CSF-edge, WM). These and the single ICA component showing the strongest correlation with motion estimates were included as additional nuisance variates. EPI volumes with excessive volume-to-volume displacement (>0.25 mm), as well as those during and immediately following the delivery of aversive reinforcers, were censored.

#### Decomposing Canonical Effects Using Finite Impulse Response (FIR) Modeling

The canonical modeling approach estimates the amplitude of anticipatory activity under the assumption that it approximates a ‘boxcar’-like square-wave function. This makes it tempting to conclude that regions showing significant activation represent sustained responses. Yet there is ample evidence that a variety of other signals are plausible (e.g. Gonzalez-Castillo et al., 2015; Gungor and Paré, 2016; Sreenivasan and D’Esposito, 2019) and, importantly, can yield similarly strong statistical effects (**Fig. 2**). Addressing this ambiguity necessitates a finer decomposition of the signal underlying significant ‘omnibus’ effects revealed by canonical modeling—a surprisingly rare approach in the neuroimaging literature. To do so, we identified the most extreme peak (e.g. BST) in each of the major regions identified in our canonical analyses. These peak locations were then interrogated using a finite impulse response (FIR) analysis, which provides an estimate of the magnitude *and* shape of anticipatory activity (Glover, 1999; Ollinger et al., 2001; for a similar approach by our group, see Guller et al., 2012). To perform the FIR modeling, variance related to reinforcer delivery, ratings, and the nuisance variables was removed from the preprocessed data using the canonical approach described above. Residualized data were bandpass filtered (.007813-0.2667 Hz) and normalized to the *SD* of the Certain Safety trials (i.e. the implicit baseline in the canonical HRF GLM). We then estimated the mean response at each TR of the anticipatory epoch for each condition of the MTC paradigm for each subject.

**Fig. 2.**
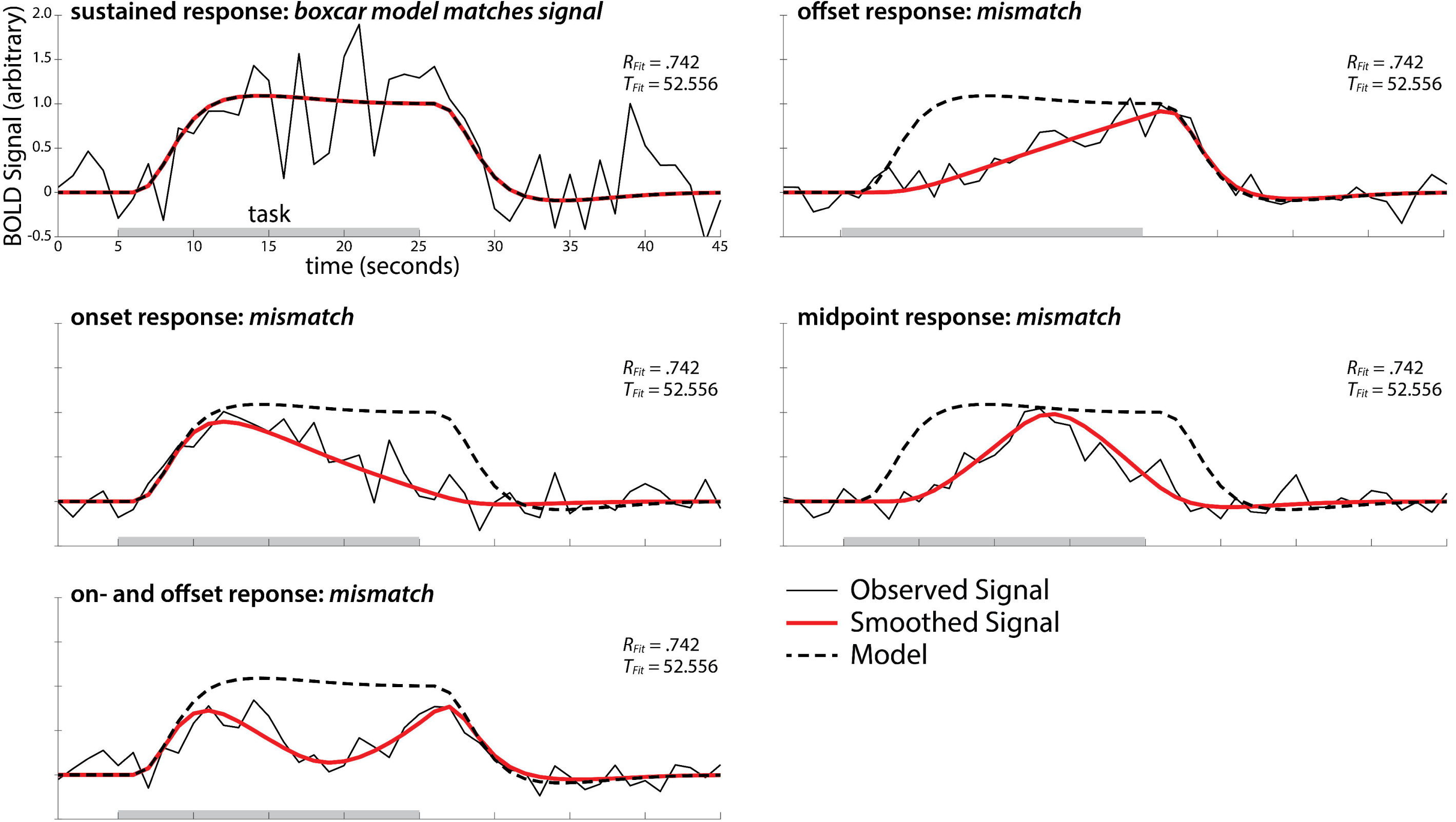
Interpretive ambiguities of canonical HRF modeling. The canonical approach to fMRI analysis models the amplitude of anticipatory activity (*solid black line*) under the assumption that it approximates a ‘boxcar’-like square-wave shape (*dotted line;* convolution of a canonical HRF with task duration). In some cases, such as the upper-left panel, the hemodynamic signal and the model will match. But in others, it will not. Importantly, a variety of physiologically plausible hemodynamic responses can produce similarly strong and statistically significant results (*T* = 52.556 in this example), highlighting the importance of modeling the BOLD signal on a finer temporal scale.

### Experimental Design and Statistical Analyses

#### Overview

Study design is described in ‘Maryland Threat Countdown (MTC) fMRI Paradigm.’ The number of usable datasets, data exclusions, and power considerations are detailed in ‘Subjects.’

#### In-Scanner Fear/Anxiety Ratings and Skin Conductance

Data were analyzed using standard repeated-measures GLM approaches with Huynh-Feldt correction for potential non-sphericity implemented in SPSS (version 24; IBM, Inc., Armonk, NY). Significant interactions were decomposed using simple effects. Figures were created using R Studio (http://www.rstudio.com) and yarrr (version 0.1.5) for R (version 3.6.1.; https://www.R-project.org).

#### Canonical Second-Level GLM

Standard whole-brain voxelwise GLMs (random effects) were used to compare anticipatory hemodynamic activity elicited by each threat-anticipation condition and its corresponding control condition (e.g. Uncertain Threat vs. Uncertain Safety). Significance was assessed using FDR *q*<.05, whole-brain corrected. As in prior work by our group (Shackman et al., 2013; Shackman et al., 2017), a minimum conjunction (logical ‘AND’) was used to identify voxels sensitive to both temporally certain *and* temporally uncertain threat anticipation (Nichols et al., 2005). We also directly examined potential differences in anticipatory activity between the two threat conditions (Certain Threat vs. Uncertain Threat). We did not examine hemodynamic responses to reinforcer delivery given the possibility of artifact. Some figures were created using MRIcron (http://people.cas.sc.edu/rorden/mricron). Clusters and local maxima were labeled using a combination of the Allen Institute, Harvard–Oxford, and Mai atlases (Frazier et al., 2005; Desikan et al., 2006b; Makris et al., 2006; Hawrylycz et al., 2012; Mai et al., 2015) and a recently established consensus nomenclature (ten Donkelaar et al., 2018).

### Descriptive Decomposition of Canonical Effects Using FIR Modeling

To decompose the signal underlying significant canonical effects, we identified the most extreme peak in each of the major regions (e.g. amygdala) identified in our canonical analyses (indicated by a black-and-white asterisk in the accompanying figures). These peaks were then descriptively interrogated using FIR models. As shown in **Fig. 1**, the duration of anticipatory epoch differed between certain (18.75 s) and uncertain trials (8.75-32.5 s; *M*=18.75 s), necessitating slightly different procedures for specific contrasts. For the comparison of Certain Threat to Certain Safety, responses were modeled for 15 TRs (1.25 s TR; total=18.125 s). Given the temporal resolution and autocorrelation of the BOLD signal, data were averaged for 3 windows (TR-1 to TR-5, TR-6 to TR-10, TR-11 to TR-15). For the comparison of Uncertain Threat to Uncertain Safety, responses were modeled for 24 TRs (total=30.00 s) and averaged for 4 windows, the first three corresponding to those used for certain trials and a fourth spanning TR-16 to TR-24. This choice was partially motivated by the modest number of trials with the longest anticipatory epoch (**Fig. 1**). For the comparison of the two threat conditions, responses were modeled for 15 TRs and averaged across 3 windows, as above. ‘Sustained’ activity was operationally defined as greater mean activity across two consecutive windows. Using this criterion, descriptive tests indicated nominally significant (*p*<.05) evidence of sustained responses for most of the key regions for most of the contrasts (e.g. Uncertain Threat vs. Uncertain Safety). Exceptions were the PAG for the Certain Threat vs. Certain Safety contrast, and the PAG and BST for the Certain Threat vs. Uncertain Threat contrast. Because this approach yields optimistically biased effect-size estimates (Kriegeskorte et al., 2010; Davenport and Nichols, 2020), we refrain from reporting exact *p*-values, and instead provide standard errors of the mean as a descriptive guide to the size of observed effects. Naturally, any inferences drawn from inspection of the standard errors only apply to the peak voxels depicted in the accompanying figures (black-and-white asterisks) and not necessarily to the entire parent region (e.g. amygdala).

#### Testing Whether the BST and Amygdala Show Different Responses to Threat

To test hypothesized regional differences in threat sensitivity (see the Introduction), we used a combination of anatomical and functional criteria to independently identify BST and amygdala voxels that were most sensitive to each kind of threat. As shown in **Fig. 3**, each region was anatomically defined using an *a priori* probabilistic region of interest (ROIs; Frazier et al., 2005; Desikan et al., 2006a; Theiss et al., 2017). Next, we extracted and averaged standardized regression coefficients for voxels showing significant (FDR *q*<.05, whole-brain corrected) activation for each of the relevant contrasts, separately for each region: Uncertain Threat > Uncertain Safety, Certain Threat > Certain Safety, and Certain Threat > Uncertain Threat. Potential regional differences (i.e. Region × Condition interactions) were then assessed using standard repeated-measures GLM approaches implemented in SPSS. Nonparametric tests (Wilcoxon signed rank) yielded identical conclusions (not reported). Inferences necessarily apply only to the subset of BST or amygdala voxels that proved sensitive to one or more of the threat contrasts, *not* the entire region.

**Fig. 3.**
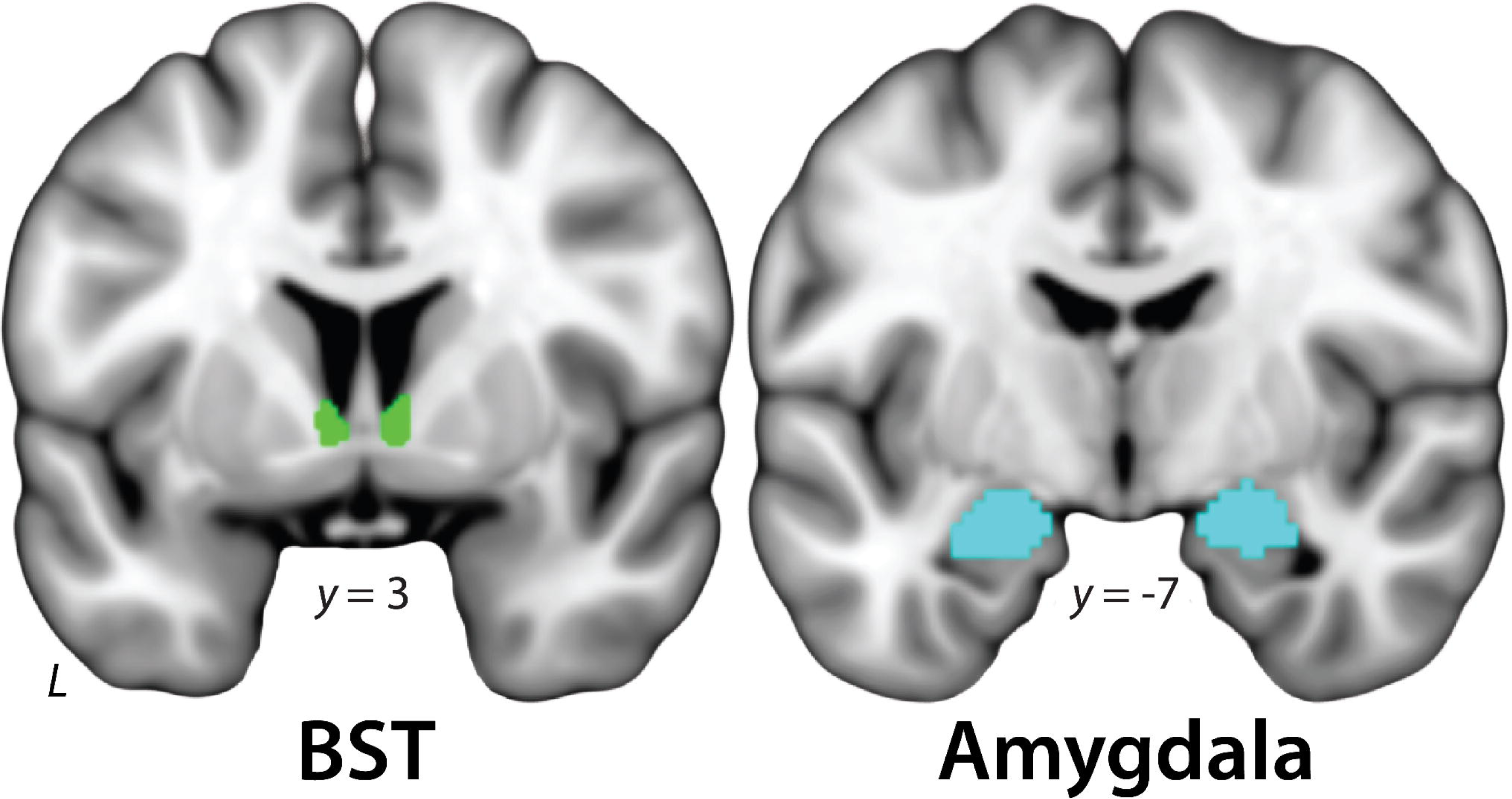
Amygdala and BST ROIs. BST. The probabilistic BST ROI (*green*) is described in (Theiss et al., 2017) and was thresholded at 0%. The seed mostly encompasses the supra-commissural BST, given the difficulty of reliably discriminating the borders of regions below the anterior commissure on the basis of T1-weighted images (Kruger et al., 2015). ***Amygdala***. The Harvard-Oxford probabilistic amygdala (*cyan*) is described in (Frazier et al., 2005; Desikan et al., 2006a) and conservatively thresholded at 50%. Analyses employed ROIs decimated to the 2-mm resolution of the EPI data. For illustrative purposes, 1-mm ROIs are shown. Single-subject data were visually inspected to ensure that the ROIs were correctly aligned to the spatially normalized T1-weighted images. Abbreviation—BST, bed nucleus of the stria terminalis.

#### Testing Whether the BST and Amygdala Show Equivalent Responses to Threat

Consistent with prior work by our group (McMenamin et al., 2009; McMenamin et al., 2010), we used the well-established two one-sided tests (TOST) procedure for formally testing whether the BST and amygdala show statistically equivalent activity during threat anticipation. While it is not possible to definitively show that the true difference in regional activity is zero, TOST provides a well-established and widely used framework for testing whether mean differences are small enough to be considered equivalent (Lakens, 2017; Lakens et al., 2018). Regression coefficients were extracted and averaged using the approach detailed in the prior section. For present purposes, we considered regional differences smaller than a ‘medium’ standardized effect for dependent means (Cohen’s *d*_z_=.35) to be equivalent (Lakens, 2017). TOST procedures were performed using the TOSTER package (version 0.3.4) for R.

## RESULTS

### Temporally Uncertain Threat anticipation elicits robust symptoms and signs of anxiety

As shown in **Fig. 1**, threat anticipation markedly increased subjective symptoms (in-scanner ratings) and objective signs (skin conductance) of anxiety, and this was particularly evident when the timing of aversive stimulation was *uncertain*. ***Anticipatory feelings***. Subjects reported experiencing more intense fear/anxiety when anticipating aversive outcomes (*F*(1,98)=543.27, *p*< .001), and when anticipating outcomes with uncertain timing (*F*(1,98)=85.46, *p*<.001). The impact of threat on fear/anxiety ratings was potentiated by temporal uncertainty (Valence × Uncertainty: *F*(1,98)=13.08, *p*<.001; Uncertain Threat > Certain Threat: *t*(98)=7.58, *p*<.001; Uncertain Safety > Certain Safety: *t*(98)=4.90, *p*<.001; Uncertain Threat > Uncertain Safety: *t*(98)=21.98, *p*<.001; Certain Threat > Certain Safety: *t*(98)=20.36, *p*<.001), consistent with prior work (Grillon et al., 2006; Nelson and Shankman, 2011; Somerville et al., 2013; Bennett et al., 2018). ***Anticipatory arousal***. Subjects showed elevated skin conductance levels when anticipating aversive outcomes (*F*(1,98)=345.31, *p*<.001), and when anticipating outcomes with uncertain timing (*F*(1,98)=85.86, *p*<.001). The impact of threat on skin conductance was potentiated by temporal uncertainty (Valence × Uncertainty: *F*(1,98)=93.63, *p*<.001; Uncertain Threat > Certain Threat: *t*(98) = 11.53, *p*<.001; Uncertain Safety > Certain Safety: *t*(98) = −3.99, *p*< .001; Uncertain Threat > Uncertain Safety: *t*(98)=25.59, *p*<.001; Certain Threat > Certain Safety: *t*(98)=9.84, *p*<.001). Taken together, these results confirm the validity of the MTC paradigm for understanding the neural circuits underpinning human anxiety.

### Temporally Uncertain Threat anticipation recruits a distributed network of subcortical and cortical regions

Next, a voxelwise GLM was used to identify brain regions recruited during the anticipation of temporally Uncertain Threat (Uncertain Threat > Uncertain Safety; FDR *q*<.05, whole-brain corrected). As shown in **Fig. 4**, this highlighted a widely distributed network of regions previously implicated in the expression and regulation of human fear and anxiety (Fullana et al., 2016; Qi et al., 2018; Fox and Shackman, 2019; Chavanne and Robinson, in press), including the MCC; AI extending into the frontal operculum (FrO); dlPFC extending to the frontal pole (FP); brainstem encompassing the periaqueductal grey (PAG); basal forebrain, in the region of the BST; and dorsal amygdala, in the region of the central and medial nuclei. Heightened activity during the anticipation of Uncertain Threat was also evident in the orbitofrontal cortex, basal ganglia, hippocampus, and ventrolateral amygdala in the region of the lateral nucleus (**Extended Data Fig. 4-1**). Consistent with prior work (Choi et al., 2012; Grupe et al., 2016), Uncertain Threat anticipation was associated with *reduced* activity in a set of midline regions that resembled the default mode network (e.g. anterior rostral sulcus/ventromedial prefrontal cortex, postcentral gyrus, and precuneus), as well as the posterior insula and parahippocampal gyrus (**Extended Data Fig. 4-2**). Reduced activity was also observed in the most rostral tip of the amygdala, underscoring the functional heterogeneity of this complex structure.

**Fig. 4.**
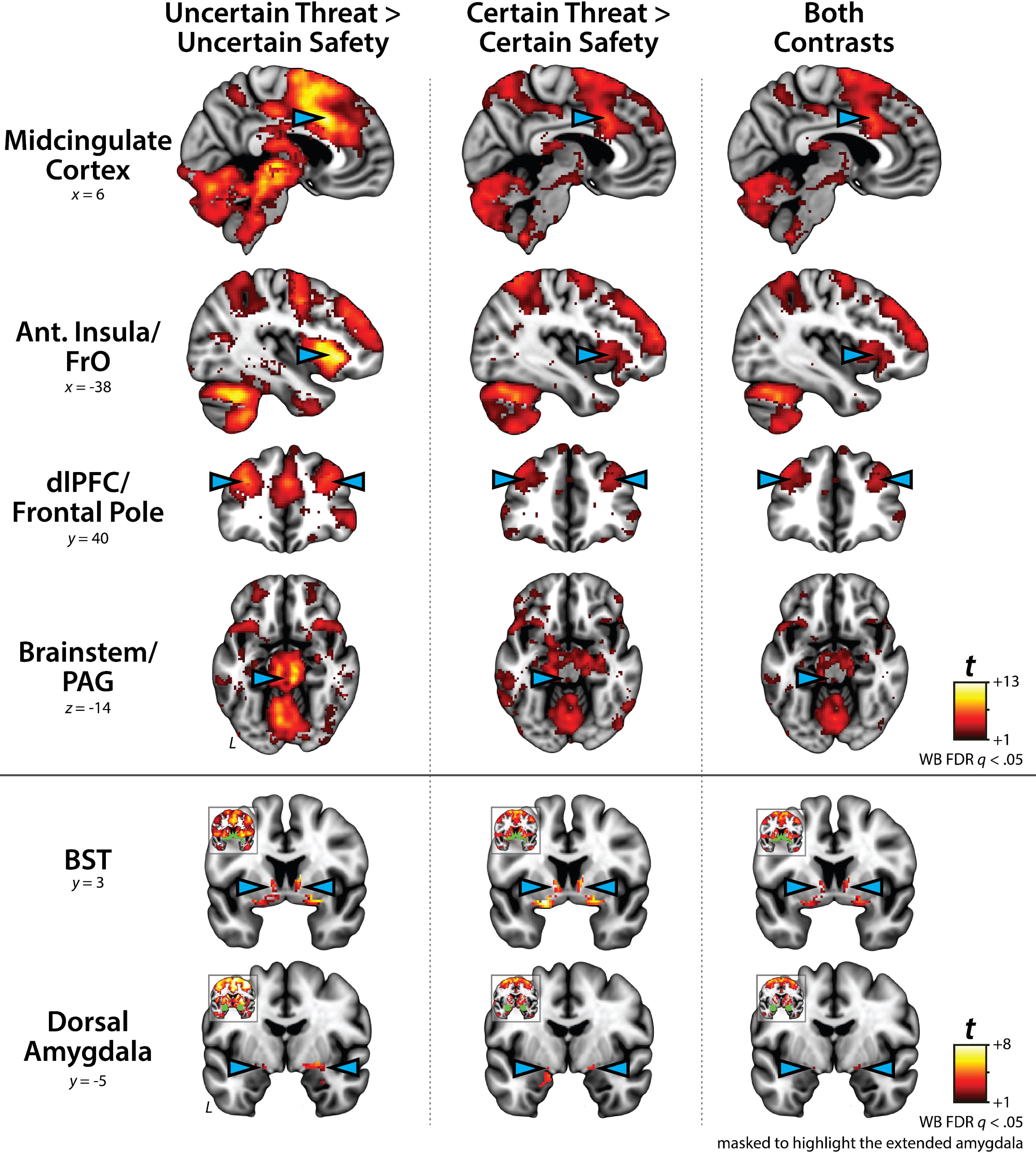
The anticipation of temporally Uncertain and Certain Threat recruits broadly similar neural systems. Key regions (*cyan arrowheads*) showing significantly elevated activity during the anticipation of Uncertain Threat (*left column*) and Certain Threat (*center column*) compared to their respective control conditions. Voxels showing significantly increased activity in both contrasts are depicted in the *right column*. BST and dorsal amygdala images are masked to highlight significant voxels in extended amygdala (*green*). Coronal insets depict the thresholded statistical parametric maps without the additional mask. Taken together, these observations indicate that these regions are sensitive to both temporally certain and uncertain threat. For additional details, see **Extended Data Figs. 4-1** to **4-5**. Abbreviations—Ant., anterior; BST, bed nucleus of the stria terminalis; dlPFC, dorsolateral prefrontal cortex; FrO, frontal operculum; L, left; PAG, periaqueductal grey; WB, whole-brain corrected.

### Temporally Uncertain Threat anticipation elicits sustained hemodynamic activity

Anxiety is widely conceptualized as a sustained state (Davis et al., 2010; Tye et al., 2011; LeDoux and Pine, 2016; Mobbs, 2018), and it is tempting to interpret clusters of enhanced activity (e.g. **Fig. 4**) through this lens. But do we actually see evidence of sustained responses during the anticipation of temporally Uncertain Threat? Although a wide variety of other signals are physiologically plausible (**Fig. 2**), the vast majority of fMRI studies never address this question; they simply assume the shape of the hemodynamic response and focus on estimates of response magnitude (‘activation’). To address this ambiguity, we used a finite impulse response (FIR) approach to estimate responses elicited by the anticipation of Uncertain Threat and Uncertain Safety on a finer time-scale. Descriptively, this revealed sustained activity (see Materials and Methods) across key cortical (MCC, AI/FrO, dlPFC/FP) and subcortical (PAG, BST, dorsal amygdala) regions (Uncertain Threat > Uncertain Safety; 6.25-30 s; **Fig. 5**).

**Fig. 5.**
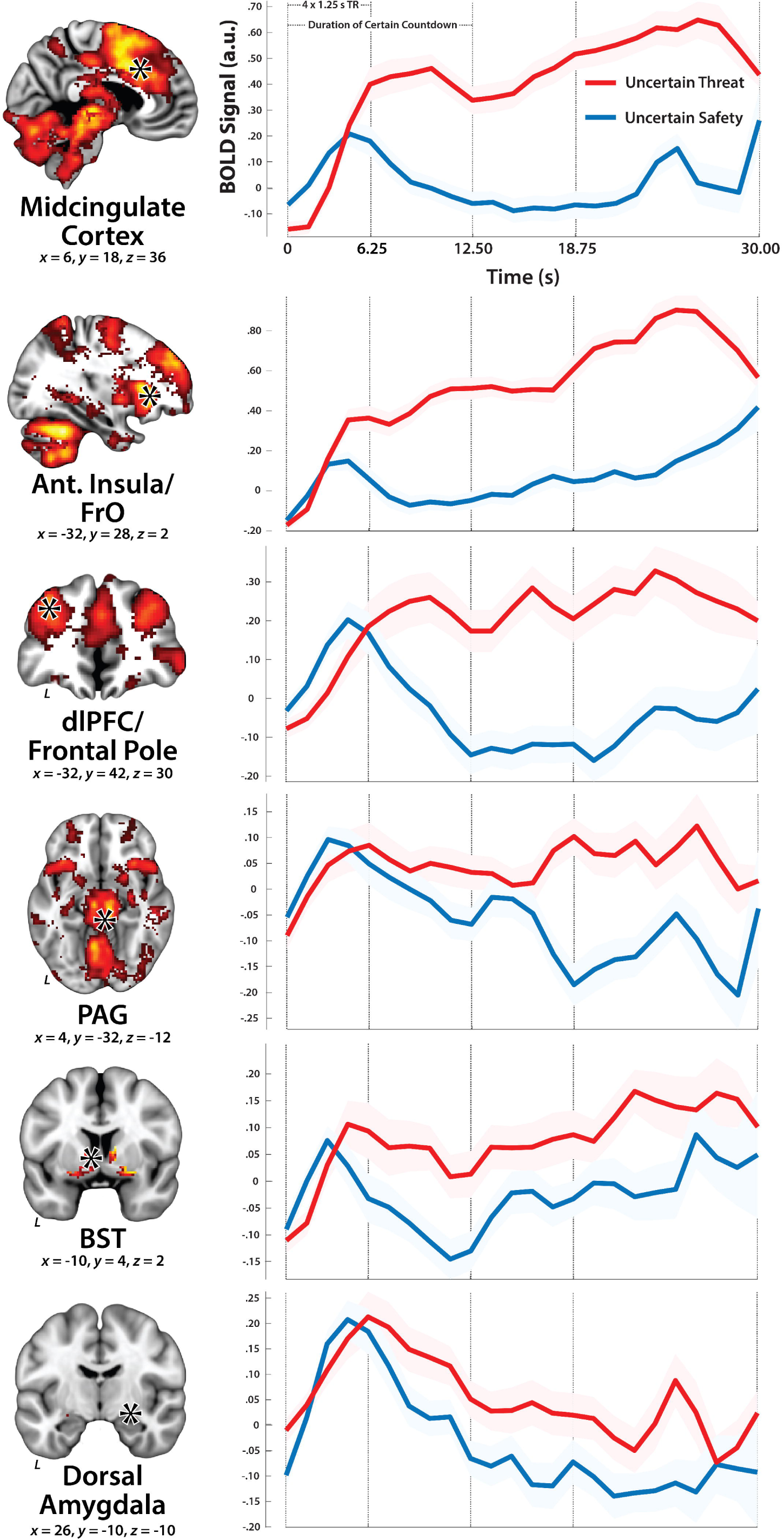
Regions sensitive to temporally Uncertain Threat show sustained hemodynamic activity. Mean responses to the anticipatory epoch were estimated on a TR-by-TR (1.25 s) basis for Uncertain Threat (*red*) and Uncertain Safety (*blue*) trials, using data from the local maxima of key clusters (*black- and-white asterisks in the left panels*) identified using a canonical analytic approach. Given the temporal resolution and autocorrelation of the hemodynamic signal, data were averaged for 4 windows (TR-1 to TR-5, TR-6 to TR-10, TR-11 to TR-15, and TR-16 to TR-24), spanning a total of 24 measurements (30 s). Windows are indicated by broken vertical lines. Shaded envelopes depict the standard error of the mean. Abbreviations—Ant., anterior; BST, bed nucleus of the stria terminalis; dlPFC, dorsolateral prefrontal cortex; FrO, frontal operculum; L, left; PAG, periaqueductal grey; TR, repetition time (the time needed to acquire a single volume of fMRI data).

### Temporally Certain Threat anticipation recruits an anatomically and functionally similar network

Having identified a distributed neural circuit sensitive to Uncertain Threat, we used a parallel approach to identify regions recruited during the anticipation of temporally *Certain* Threat (Certain Threat > Certain Safety; FDR *q*<.05, whole-brain corrected). As shown in **Fig. 4**, results were similar to those found for Uncertain Threat (**Extended Data Figs. 4-3** and **4-4**). In fact, a minimum conjunction analysis (Logical ‘AND;’ Nichols et al., 2005) revealed voxelwise co-localization in every key cortical and subcortical region, including the BST and dorsal amygdala in the region of the central and medial nuclei (**Fig. 4** and **Extended Data Fig. 4-5**). FIR results also suggested functional convergence across conditions, with all but one of these key regions (PAG) showing sustained levels of heightened hemodynamic activity during the anticipation of Certain Threat (see Materials and Methods; **Fig. 6**). Taken together, these results suggest that this network of subcortical and cortical regions is sensitive to multiple kinds of threat anticipation, both certain and uncertain.

**Fig. 6.**
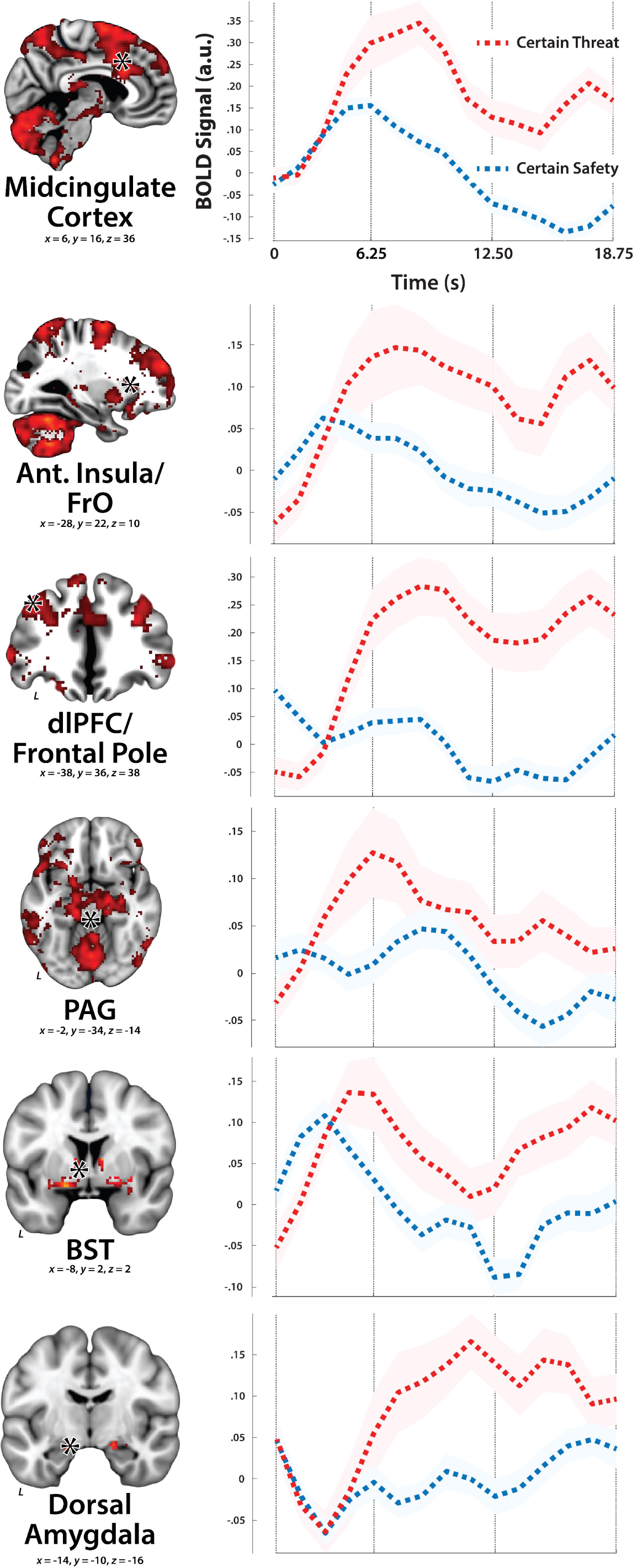
Regions sensitive to temporally Certain Threat show sustained hemodynamic activity. Mean responses to the anticipatory epoch were estimated on a TR-by-TR (1.25 s) basis for Certain Threat (*red*) and Certain Safety (*blue*) trials, using data from the local maxima of key clusters (*black-and-white asterisks in the left panels*) identified using a canonical HRF GLM approach. Given the temporal resolution and autocorrelation of the hemodynamic signal, data were averaged for 3 windows (TR-1 to TR-5, TR-6 to TR-10, and TR-11 to TR-15), spanning a total of 15 measurements (18.75 s). Windows are indicated by broken vertical lines. Shaded envelopes depict the standard error of the mean. Abbreviations—Ant., anterior; BST, bed nucleus of the stria terminalis; dlPFC, dorsolateral prefrontal cortex; FrO, frontal operculum; L, left; PAG, periaqueductal grey.

### The threat anticipation network can be fractionated into subdivisions

To determine whether regions recruited during threat anticipation are sensitive to temporal uncertainty, we directly compared the Uncertain and Certain Threat conditions (FDR *q*<.05, whole-brain corrected). This indicated that the threat anticipation network can be fractionated into subdivisions. As shown in **Fig. 7**, key cortical regions (MCC, AI/FrO, and dlPFC/FP) showed a further increase in activity during the anticipation of *Uncertain* Threat (**Extended Data Fig. 7-1**). In contrast, the BST and dorsal amygdala (adjacent to the central nucleus, in the region of the cortical nucleus and Amygdala-Hippocampal Transition Area) showed the reverse pattern, with relatively greater activity during the anticipation of *Certain* Threat (**Extended Data Fig. 7-2**). The PAG did not discriminate the two threat conditions. FIR results suggest a similar conclusion (**Fig. 7**).

**Fig. 7.**
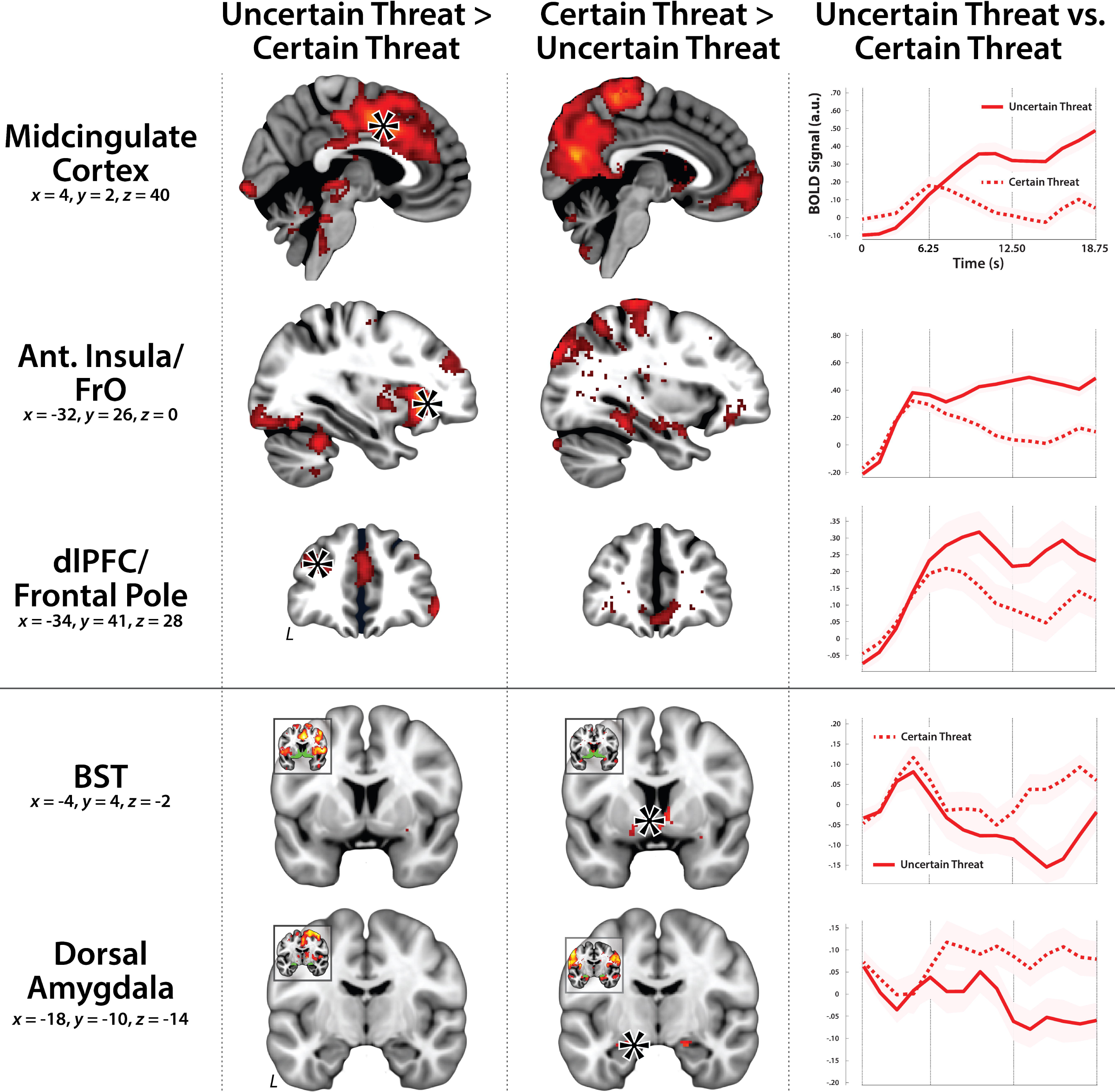
The threat anticipation network can be fractionated into subdivisions. The midcingulate cortex, anterior insula/frontal operculum, and dlPFC showed greater activity during the anticipation of Uncertain Threat (*left column*), whereas the BST and dorsal amygdala showed greater activity during the anticipation of Certain Threat (*center column*). Thresholds and other conventions are identical to **Fig. 4**. For additional details, see **Extended Data Figs. 7-1** and **7-2**. The r*ight column* depicts TR-by-TR (1.25 s) hemodynamic responses during the anticipation of Uncertain Threat (*solid red line*) and Certain Threat (*broken red line*). Data were extracted from the local maxima of key clusters (*black-and-white asterisks in the left and center columns*) identified using a canonical HRF GLM approach. Given the temporal resolution and autocorrelation of the hemodynamic signal, data were averaged for 3 windows (TR-1 to TR-5, TR-6 to TR-10, and TR-11 to TR-15), spanning a total of 15 measurements (18.75 s). Windows are indicated by broken vertical lines. Shaded envelopes depict the standard error of the mean. Abbreviations—Ant., anterior; BST, bed nucleus of the stria terminalis; dlPFC, dorsolateral prefrontal cortex; FrO, frontal operculum; L, left; PAG, periaqueductal grey.

### The anticipation of temporally Uncertain and Certain Threat elicits statistically indistinguishable responses in the extended amygdala

Our results indicate that the BST and dorsal amygdala—the two major subdivisions of the EA—respond similarly to threat anticipation. Both regions show signs of elevated activity during threat anticipation, and this is evident whether or not the timing of aversive stimulation is uncertain (**Fig. 4**). Furthermore, both regions showed parallel increases in activity during the anticipation of Certain Threat (**Figs. 6-7**). Yet it remains possible that the BST and the amygdala exhibit subtler differences in threat sensitivity. To rigorously test this, we directly compared regional responses for each of the threat contrasts (e.g. Uncertain Threat vs. Uncertain Safety), equivalent to testing the Region × Condition interactions. As shown in **Fig. 8**, mean differences were small to very-small (*d*_Z_<.17) and all non-significant (**Extended Data Fig. 8-1**). Likewise, the proportion of subjects showing numerically greater activity in one region or the other never exceeded 55% (**Fig. 8**). Naturally, these results do not license strong claims of regional equivalence. While it is impossible to demonstrate that the true difference in regional activity is zero, the two one-sided tests (TOST) procedure provides a well-established and widely used framework for testing whether mean differences—here, in regional activity—are small enough to be considered statistically equivalent (Lakens, 2017; Lakens et al., 2018). For present purposes, we considered differences smaller than a ‘medium’ standardized effect (Cohen’s *d*_z_=.35) to be statistically equivalent. Using the voxels that were most sensitive to each threat contrast (see Materials and Methods), our results revealed significant equivalence for all three contrasts (*p*s=.001-.03; **Fig. 8** and **Extended Data Fig. 8-1**). Although these statistical findings do not demonstrate that the amygdala and the BST are functionally interchangeable (‘the same’), they do enable us to decisively reject claims of strict functional segregation (i.e. that the BST is sensitive to uncertain danger, whereas the amygdala is not) for the subset of these regions engaged by the MTC paradigm.

**Fig. 8.**
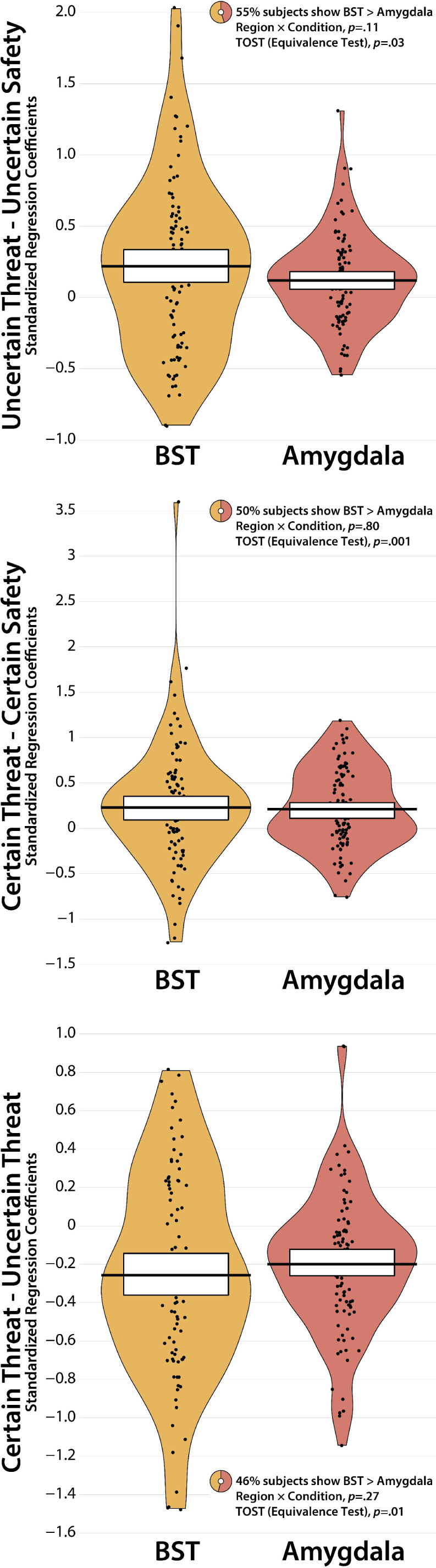
The BST and dorsal amygdala regions recruited by the Maryland Threat Countdown paradigm show statistically indistinguishable responses during threat anticipation. While it is impossible to demonstrate that the true difference in regional hemodynamic activity is zero, the two one-sided tests (TOST) procedure provides a well-established and widely used statistical framework for testing whether mean differences in regional activity are small enough to be considered equivalent (Lakens, 2017; Lakens et al., 2018). Using the subset of voxels that were most sensitive to each threat contrast (see Materials and Methods), results revealed significant equivalence for all contrasts (**Extended Data Fig. 8-1**). Figure depicts the data (*black points; individual participants*), density distribution (*bean plots*), Bayesian 95% highest density interval (HDI; *colored bands*), and mean (*black bars*) for each condition. HDIs permit population-generalizable visual inferences about mean differences and were estimated using 1,000 samples from a posterior Gaussian distribution. Inset ring plots indicate the percentage of subjects showing greater activity in the BST compared to the dorsal amygdala for each contrast.

## DISCUSSION

Uncertain-threat anticipation is the prototypical trigger of anxiety, a core theme that cuts across psychiatric disorders, species, and assays, including novelty, darkness, and other ‘diffuse’ threats. Despite the immense significance of anxiety for public health, the neural systems recruited by uncertain threat have remained unclear. Leveraging a translationally relevant paradigm optimized for fMRI signal decomposition (**Fig. 1**), our results reveal that the anticipation of temporally uncertain aversive stimulation recruits a distributed network of fronto-cortical (MCC, AI/FrO, and dlPFC/FP) and subcortical (PAG, BST, and dorsal amygdala) regions (**Fig. 4**), mirroring robust changes in experience and psychophysiology (**Fig. 1**). Closer inspection of signal dynamics in these regions provided descriptive support for sustained activity during the anticipation of Uncertain Threat (**Fig. 5**). Analyses focused on the anticipation of temporally Certain Threat revealed a similar pattern, with voxels sensitive to both kinds of threat evident in key cortical and subcortical regions (**Fig. 4**), suggesting that this circuitry is sensitive to both certain and uncertain threat. Direct comparison of the two threat conditions demonstrated that this network can be fractionated: cortical regions showed relatively greater activity during the anticipation of Uncertain Threat, whereas the extended amygdala showed relatively greater activity during the anticipation of Certain Threat (**Fig. 7**). While there is consensus that the BST and dorsal amygdala play a critical role in orchestrating adaptive responses to danger, their precise contributions to human anxiety have remained contentious. Our results suggest that these regions respond similarly to different kinds of threat anticipation. In fact, we show that the BST and dorsal amygdala exhibit statistically indistinguishable responses to threat anticipation across a variety of comparisons (**Fig. 8**), reinforcing the possibility that they make broadly similar contributions to human anxiety (Gungor and Paré, 2016; Fox and Shackman, 2019).

Since the time of Freud (Freud, 1920), the distinction between certain (*‘fear’*) and uncertain (*‘anxiety’*) danger has been a key feature of neuropsychiatric models of emotion (Davis et al., 2010; LeDoux and Pine, 2016; Mobbs, 2018). Our findings show that the regions recruited during the anticipation of Certain and Uncertain Threat are co-localized in several key regions (**Fig. 4**). This common threat anticipation network encompasses subcortical regions that are critical for assembling defensive responses to uncertain threat in animals (Fox and Shackman, 2019). But it also includes fronto-cortical regions—like the MCC, AI/FrO, and dlPFC/FP—that have received less empirical attention and are challenging to study in rodents (e.g. Carlén, 2017). These regions have traditionally been associated with the controlled processing and regulation of emotion and cognition (Shackman et al., 2011; Morawetz et al., 2017; Langner et al., 2018; Kroes et al., 2019; Picó-Pérez et al., 2019; Morawetz et al., *in press*) and more recently implicated in the conscious experience of emotion (LeDoux, 2020). As shown in **Fig. 9**, the present results are well aligned with recent meta-analyses of neuroimaging studies of ‘fear’ (Fullana et al., 2016) and ‘anxiety’ (Chavanne and Robinson, in press). Across studies encompassing tens of studies and hundreds of subjects, this work demonstrates that the anticipation of *certain-*threat (Pavlovian threat cues; the prototypical ‘fear’ stimulus in laboratory studies) and *uncertain-*threat (instructed ‘threat-of-shock’) recruit an overlapping network of core regions, including the BST (but not the Ce; see below). This similarity cannot be dismissed as an artifact of neuroimagers’ penchant for partial-reinforcement Pavlovian paradigms, which render ‘certain’ threat uncertain (Fullana et al., 2016, median threat probability=63%; Picó-Pérez et al., 2019, median threat probability=62%). In fact, the same general pattern—including elevated activity in the region of the BST—is evident in large-scale studies of *certain-* (Sjouwerman et al., 2020, Study 2, n=113, threat probability=100%, https://neurovault.org/collections/6031) and *uncertain*-threat anticipation (Klumpers et al., 2017, Sample 1: n=108, threat probability=33%), consistent with our results.

**Fig. 9.**
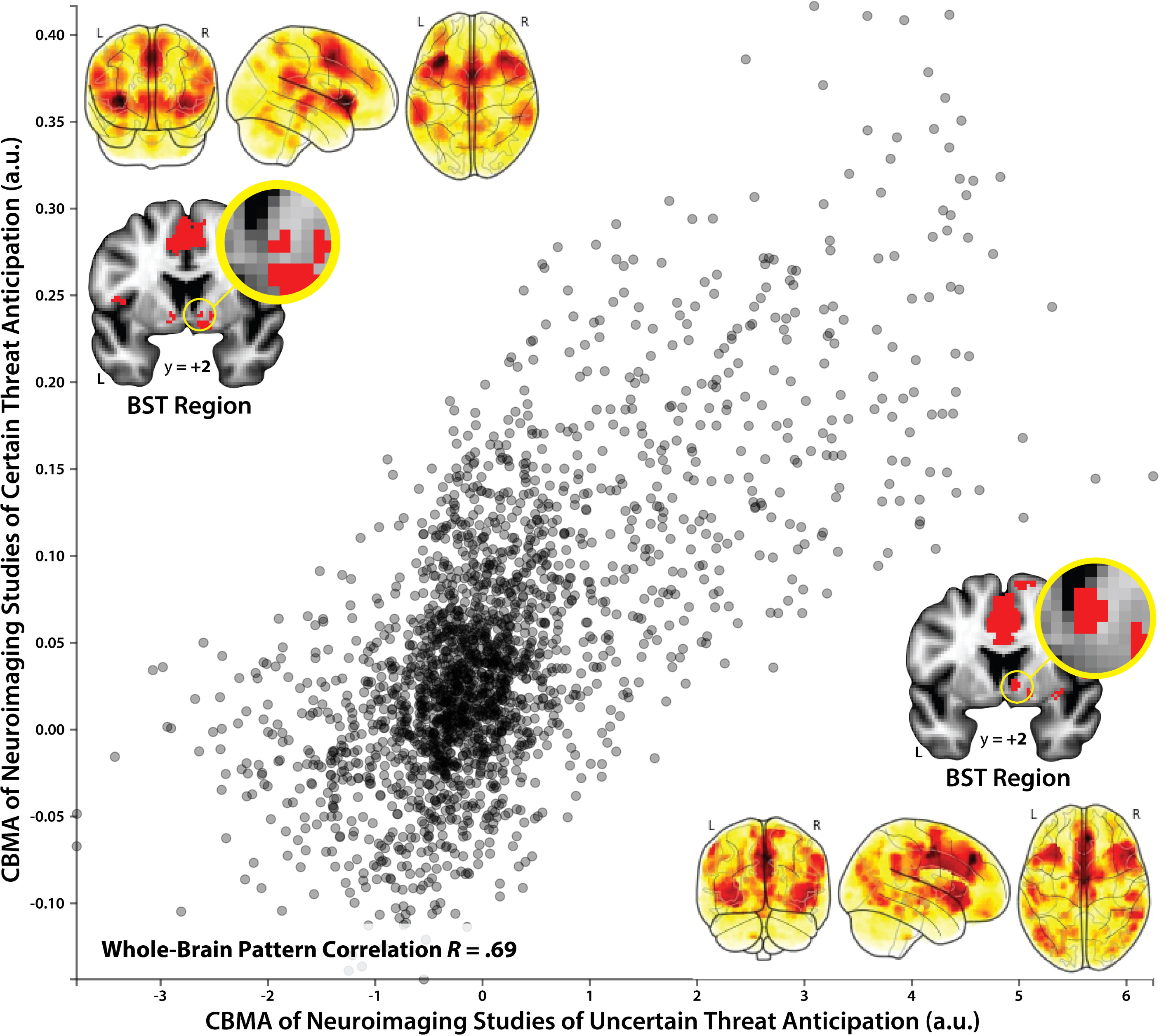
Certain and uncertain threat anticipation elicit broadly similar patterns of neural activity. Figure summarizes the results of two coordinate-based meta-analyses (CBMA) of functional neuroimaging studies. *Top-left* inset depicts the results for 27 ‘fear conditioning’ studies (*N*=677), highlighting regions showing consistently greater activity during the anticipation of certain threat (CS+ > CS-; https://neurovault.org/collections/2472; Fullana et al., 2016). *Bottom-right* inset depicts the results for 18 ‘threat-of-shock’ studies (*N*=693), highlighting regions showing consistently greater activity during the anticipation of uncertain threat (Threat > Safe; https://neurovault.org/collections/6012; Chavanne and Robinson, in press). Visual inspection of the results (*red clusters*) suggests that the anticipation of certain and uncertain threat elicits qualitatively similar patterns, including heightened activity in the region of the BST. This impression is reinforced by the substantial correlation between the two whole-brain patterns, *r* = .69. Consistent amygdala activity was not detected in either meta-analysis. Note: The pattern correlation was estimated in Neurovault using a brain-masked, 4-mm transformation of the publicly available, vectorized meta-analytic maps (Gorgolewski et al., 2015). For illustrative purposes, every 10^th^ voxel is depicted in the scatter plot. Abbreviations—BST, bed nucleus of the stria terminalis; CBMA, coordinate-based meta-analyses; L, left hemisphere; R, right hemisphere.

Our observations provide insight into the functional architecture of the threat anticipation network, demonstrating that fronto-cortical regions prefer Uncertain over Certain Threat, whereas the BST and dorsal amygdala show the reverse preference—*a difference in degree, not in kind*. Trivial differences cannot account for this nuance; the two threat conditions were pseudo-randomly intermixed and nearly identical in terms of their perceptual, nociceptive, motor, and statistical features (**Fig. 1**). What might explain the observed regional preferences? Aside from temporal certainty, the most conspicuous difference between the conditions is the degree of cognitive scaffolding. On Certain Threat trials, the descending stream of integers provided a precise and predictable index of momentary changes in threat imminence, encouraging a reactive, stimulus-bound cognitive mode. On Uncertain Threat trials this support was absent, necessitating greater reliance on the kinds of sustained, endogenous representations that are the hallmark of fronto-cortical regions (Badre and Nee, 2018). A second notable difference between the two threat conditions is the intensity of anxiety. Uncertain-Threat anticipation was associated with greater distress and arousal (**Fig. 1**). The observed increase in fronto-cortical activity could reflect either heightened anxiety or compensatory processes aimed at downregulating distress and arousal. Testing these non-exclusive hypotheses will require a multi-pronged approach that encompasses carefully optimized tasks, mechanistic interventions, and a broader assessment of the nomological network. Multivoxel classifier approaches are likely to be useful for linking specific facets of anxiety (e.g. feelings) to particular elements of the threat anticipation network, and determining whether this reflects expressive or regulatory processes (Chang et al., 2015).

The present results add to a growing body of evidence indicating that the BST and dorsal amygdala, while certainly not interchangeable, are more alike than different (Fox and Shackman, 2019). The BST and dorsal amygdala are characterized by broadly similar patterns of anatomical connectivity, cellular composition, neurochemistry, and gene expression (Fox et al., 2015), although some differences in functional connectivity have been identified (Gorka et al., 2018). Both regions are poised to trigger defensive responses via dense projections to downstream effectors (Fox et al., 2015). Neuroimaging studies have documented similar responses in the two regions to a range of anxiety-eliciting stimuli (Fox and Shackman, 2019; Hudson et al., 2020, https://neurovault.org/collections/6237), and mechanistic work in rodents reinforces the hypothesis that the BST and dorsal amygdala (Ce) are crucial substrates for human anxiety (Fox and Shackman, 2019). In fact, work using a variant of the present paradigm in mice shows that Ce-BST projections are necessary for mounting defensive responses during the anticipation of temporally uncertain shock (Lange et al., 2017), consistent with our general conclusions. While our understanding remains far from complete, this body of observations underscores the need to revise models of anxiety, like RDoC, that imply a strict segregation of certain and uncertain threat processing in the extended amygdala. The present results imply that the magnitude of regional differences in hemodynamic sensitivity to threat-uncertainty is modest (<*dz*=.35); conditional on perceptual confounds, collinearities, or other moderators; or simply non-existent. An important challenge for the future will be to determine whether the *type* of threat uncertainty (e.g. temporal vs. likelihood) is a crucial determinant of regional differences in function.

Our results indicate that the amygdala’s response to threat anticipation is sparse, at least when compared to widely used emotional face and scene paradigms. This was not unexpected. The amygdala is a heterogeneous collection of at least 13 nuclei and cortical areas—*not* ‘a thing’ (Swanson and Petrovich, 1998; Yilmazer-Hanke, 2012)—and converging lines of mechanistic and imaging evidence point to the special importance of the dorsal amygdala, in the region of the Ce (Davis et al., 2010; Fox and Shackman, 2019; Hur et al., 2019). In humans, Ce represents ∼3% of total amygdala volume (Wegiel et al., 2014; Avino et al., 2018). The dorsal amygdala clusters that we observed extend beyond the Ce to encompass neighboring dorso-caudal aspects of the medial, lateral, and cortical nuclei, and amygdala-hippocampal transition area (Extended Data). While meta-analyses of small-sample neuroimaging studies have failed to detect significant amygdala responses to threat anticipation (Fullana et al., 2016, median n=16; Chavanne and Robinson, in press, median n=29), the location and extent of the dorsal amygdala clusters reported here align with more recent large-sample studies of *certain-* (Sjouwerman et al., 2020) and *uncertain*-threat anticipation (Reddan et al., 2018, n=68, threat probability=33%). In sum, our results are broadly aligned with amygdala anatomy, prior theory, and emerging neuroimaging evidence.

To conclude, the neural circuits recruited by temporally uncertain and certain threat are not categorically different, at least when viewed through the macroscopic lens of fMRI. We see evidence of anatomical co-localization—*not* segregation—in a number of key regions, in broad accord with animal models and recent imaging meta-analyses. This shared threat-anticipation system can be fractionated, with fronto-cortical regions showing relatively stronger engagement during the anticipation of temporally uncertain threat, and the BST and dorsal amygdala showing the reverse pattern. In direct comparisons, the BST and dorsal amygdala exhibited statistically indistinguishable responses, reinforcing the possibility that they make similar contributions to human anxiety. These observations provide a framework for conceptualizing fear and anxiety and for guiding mechanistic work aimed at developing more effective intervention strategies for pathological anxiety. A large sample, well-controlled task, and advanced techniques for data acquisition and processing enhance confidence in the robustness and translational relevance of these results.

## ACKNOWLEDGEMENTS

Authors acknowledge assistance and critical feedback from A. Antonacci, M. Barstead, D. Bradford, J. Curtin, L. Friedman, J. Furcolo, C. Gorgolewski, C. Grubb, D. Holley, R. Hum, C. Kaplan, C. Lejuez, D. Limon, B. Nacewicz, S. Padmala, L. Pessoa, M. Roesch, S. Rose, J. Swayambunathan, A. Vogel, members of the Affective and Translational Neuroscience laboratory, the staff of the Maryland Neuroimaging Center, the Office of the Registrar (University of Maryland), and two anonymous reviewers. This work was supported by the California National Primate Center; National Institutes of Health (DA040717, MH107444, MH121409); University of California, Davis; and University of Maryland, College Park. Authors declare no conflicts of interest.

