## Supplement for "Anxiety and the neurobiology of temporally uncertain threat anticipation"

Juyoen Hur^1^*

Jason F. Smith^2^*

Kathryn A. DeYoung^2,3^*

Allegra S. Anderson^6^

Jinyi Kuang^7^

Hyung Cho Kim^2,4^

Rachael M. Tillman^2^

Manuel Kuhn^8^

Andrew S. Fox^9,10^

Alexander J. Shackman^2,4,5^

^1^Department of Psychology, Yonsei University, Seoul 03722, Republic of Korea. Departments of ^2^Psychology, and ^3^Family Science; ^4^Neuroscience and Cognitive Science Program; and ^5^Maryland Neuroimaging Center, University of Maryland, College Park, MD 20742 USA. ^6^Department of Psychological Sciences, Vanderbilt University, Nashville, TN 37240 USA. ^7^Department of Psychology, University of Pennsylvania, PA 19104 USA. ^8^Center for Depression, Anxiety and Stress Research, McLean Hospital, Harvard Medical School, Belmont, MA 02478 USA. ^9^Department of Psychology and ^10^California National Primate Research Center, University of California, Davis, CA 95616 USA

* contributed equally

**Address Correspondence to:**

Juyoen Hur^1^ or Alexander J. Shackman^2,4,5^

**EXTENDED DATA FIG. 4-1**

**Descriptive statistics for clusters and local extrema showing greater activity during the anticipation of Uncertain Threat compared to Uncertain Safety (FDR *q*<.05, whole-brain corrected).**

|  | | **mm^3^** | ***t*** | ***x*** | ***y*** | ***z*** |
| --- | --- | --- | --- | --- | --- | --- |
|  | Cluster 1 | 685,144 |  |  |  |  |
|  | L Anterior Orbital Gyrus |  | 4.26 | -26 | 50 | -14 |
|  | R Anterior Orbital Gyrus |  | 3.19 | 26 | 46 | -14 |
|  | R Frontal Pole |  | 7.59 | 28 | 46 | 26 |
|  | L Middle Frontal Gyrus |  | 6.92 | -36 | 34 | 38 |
|  | L Paracingulate Gyrus |  | 8.13 | 0 | 34 | 30 |
|  | R Paracingulate Gyrus |  | 8.33 | 6 | 32 | 30 |
|  | R Frontal Orbital Cortex |  | 11.39 | 32 | 30 | 2 |
|  | L Frontal Orbital Cortex |  | 13.68 | -34 | 26 | 0 |
|  | R Inferior Frontal Gyrus, pars triangularis |  | 5.20 | 54 | 24 | 20 |
|  | L Cingulate Gyrus, anterior division |  | 10.49 | -4 | 20 | 34 |
|  | R Cingulate Gyrus, anterior division |  | 11.95 | 6 | 18 | 36 |
|  | R Frontal Operculum Cortex |  | 13.29 | 48 | 16 | 2 |
|  | L Frontal Operculum Cortex |  | 11.56 | -42 | 14 | 2 |
|  | R Inferior Frontal Gyrus, pars opercularis |  | 6.58 | 38 | 12 | 26 |
|  | L Inferior Frontal Gyrus, pars opercularis |  | 10.00 | -54 | 10 | 2 |
|  | R Juxtapositional Lobule Cortex |  | 10.97 | 4 | 8 | 56 |
|  | R Superior Frontal Gyrus |  | 10.18 | 14 | 8 | 64 |
|  | R Central Opercular Cortex |  | 11.24 | 50 | 8 | 2 |
|  | R Temporal Pole |  | 6.51 | 44 | 6 | -38 |
|  | L Temporal Pole |  | 4.64 | -40 | 4 | -42 |
|  | L Superior Frontal Gyrus |  | 9.23 | -16 | 4 | 66 |
|  | L Accumbens |  | 3.68 | -10 | 4 | -6 |
|  | R Putamen |  | 10.89 | 30 | 4 | -2 |
|  | L Bed Nucleus of the Stria Terminalis |  | 4.59 | -10 | 4 | 2 |
|  | L Central Opercular Cortex |  | 8.73 | -50 | 2 | 4 |
|  | L Putamen |  | 9.05 | -28 | 2 | -2 |
|  | R Pallidum |  | 8.38 | 16 | 2 | 0 |
|  | R Precentral Gyrus |  | 10.92 | 46 | 2 | 50 |
|  | L Inferior Temporal Gyrus, anterior division |  | 5.12 | -44 | 0 | -34 |
|  | L Insular Cortex |  | 5.05 | -32 | 0 | 14 |
|  | L Precentral Gyrus |  | 7.56 | -42 | -2 | 44 |
|  | L Parahippocampal Gyrus, anterior division |  | 3.21 | -32 | -2 | -32 |
|  | L Caudate |  | 7.62 | -16 | -2 | 20 |
|  | R Caudate |  | 8.87 | 16 | -2 | 18 |
|  | R Amygdala (*in the region of the lateral nucleus*)^a,b^ |  | 4.39 | 30 | -2 | -26 |
|  | R Middle Frontal Gyrus |  | 7.52 | 32 | -2 | 44 |
|  | R Planum Polare |  | 4.36 | 42 | -2 | -18 |
|  | L Pallidum |  | 6.19 | -14 | -4 | 0 |
|  | L Temporal Fusiform Cortex, anterior division |  | 4.38 | -32 | -6 | -38 |
|  | L Juxtapositional Lobule Cortex |  | 6.43 | -8 | -6 | 60 |
|  | R Insular Cortex |  | 4.32 | 44 | -6 | -4 |
|  | R Amygdala (*in the region of the central nucleus*)^a,c^ |  | 4.85 | 26 | -10 | -10 |
|  | L Planum Polare |  | 3.48 | -44 | -10 | -12 |
|  | R Parahippocampal Gyrus, anterior division |  | 3.38 | 20 | -12 | -32 |
|  | R Hippocampus |  | 5.12 | 30 | -14 | -12 |
|  | R Inferior Temporal Gyrus, posterior division |  | 4.39 | 54 | -16 | -32 |
|  | L Thalamus |  | 6.75 | -4 | -20 | -2 |
|  | R Thalamus |  | 10.10 | 6 | -20 | -2 |
|  | R Temporal Fusiform Cortex, posterior division |  | 2.18 | 36 | -20 | -30 |
|  | L Postcentral Gyrus |  | 6.13 | -56 | -24 | 24 |
|  | L Cingulate Gyrus, posterior division |  | 8.51 | -8 | -24 | 40 |
|  | R Cingulate Gyrus, posterior division |  | 8.00 | 10 | -24 | 40 |
|  | L Supramarginal Gyrus, anterior division |  | 6.30 | -60 | -26 | 22 |
|  | L Hippocampus |  | 3.66 | -34 | -26 | -10 |
|  | R Supramarginal Gyrus, anterior division |  | 8.42 | 56 | -26 | 26 |
|  | R Cingulate Gyrus, posterior division |  | 8.00 | 10 | -24 | 40 |
|  | L Supramarginal Gyrus, anterior division |  | 6.30 | -60 | -26 | 22 |
|  | L Hippocampus |  | 3.66 | -34 | -26 | -10 |
|  | R Supramarginal Gyrus, anterior division |  | 8.42 | 56 | -26 | 26 |
|  | L Temporal Fusiform Cortex, posterior division |  | 3.71 | -38 | -30 | -24 |
|  | L Parahippocampal Gyrus, posterior division |  | 3.63 | -26 | -30 | -28 |
|  | L Superior Temporal Gyrus, posterior division |  | 3.51 | -58 | -32 | 4 |
|  | R Middle Temporal Gyrus, posterior division |  | 8.34 | 52 | -32 | -6 |
|  | L Brainstem/Periaqueductal Gray |  | 6.23 | 4 | -32 | -12 |
|  | L Inferior Temporal Gyrus, posterior division |  | 4.12 | -56 | -34 | -26 |
|  | L Middle Temporal Gyrus, posterior division |  | 4.62 | -54 | -34 | -8 |
|  | R Postcentral Gyrus |  | 5.78 | 42 | -34 | 54 |
|  | L Parietal Operculum Cortex |  | 7.45 | -58 | -36 | 26 |
|  | R Parietal Operculum Cortex |  | 2.50 | 42 | -36 | 22 |
|  | R Supramarginal Gyrus, posterior division |  | 9.66 | 56 | -42 | 32 |
|  | L Supramarginal Gyrus, posterior division |  | 8.73 | -62 | -44 | 32 |
|  | R Middle Temporal Gyrus, temporooccipital part |  | 7.68 | 58 | -44 | 6 |
|  | R Superior Parietal Lobule |  | 7.51 | 36 | -46 | 64 |
|  | R Inferior Temporal Gyrus, temporooccipital part |  | 4.19 | 56 | -46 | -14 |
|  | L Temporal Occipital Fusiform Cortex |  | 2.67 | -36 | -48 | -12 |
|  | L Superior Parietal Lobule |  | 6.98 | -30 | -50 | 66 |
|  | R Precuneus Cortex |  | 5.11 | 10 | -50 | 52 |
|  | L Angular Gyrus |  | 5.71 | -60 | -54 | 24 |
|  | L Precuneus Cortex |  | 5.42 | -10 | -54 | 56 |
|  | L Middle Temporal Gyrus, temporooccipital part |  | 6.37 | -48 | -60 | 8 |
|  | L Lateral Occipital Cortex, superior division |  | 5.18 | -16 | -60 | 52 |
|  | R Temporal Occipital Fusiform Cortex |  | 2.70 | 42 | -60 | -14 |
|  | L Lateral Occipital Cortex, inferior division |  | 4.74 | -38 | -64 | 14 |
|  | R Lateral Occipital Cortex, inferior division |  | 4.32 | 44 | -64 | 10 |
|  | R Angular Gyrus |  | 3.63 | -18 | -66 | 48 |
|  | R Lateral Occipital Cortex, inferior division |  | 4.55 | 42 | -70 | -10 |
|  | R Occipital Fusiform Gyrus |  | 6.51 | 20 | -76 | -24 |
|  | R Lingual Gyrus |  | 5.62 | 6 | -82 | -10 |
|  | L Cuneal Cortex |  | 3.48 | -8 | -84 | 28 |
|  | L Lingual Gyrus |  | 4.45 | -12 | -86 | -10 |
|  | L Occipital Fusiform Gyrus |  | 3.27 | -12 | -88 | -16 |
|  | R Occipital Pole |  | 5.40 | 26 | -94 | 16 |
|  | L Occipital Pole |  | 3.96 | -14 | -100 | 12 |
|  | Cluster 2 | 368 |  |  |  |  |
|  | L Subcallosal Cortex |  | 3.05 | -2 | 26 | 0 |
|  | R Subcallosal Cortex |  | 4.60 | 0 | 22 | -6 |
|  | Cluster 3 | 72 |  |  |  |  |
|  | L Inferior Temporal Gyrus, temporooccipital part |  | 2.48 | -60 | -56 | -14 |
|  | Cluster 4 | 32 |  |  |  |  |
|  | R Superior Temporal Gyrus, anterior division |  | 2.48 | 54 | 4 | -12 |

^a^ Within the Harvard-Oxford amygdala (*p*>.25), a total of 72 voxels (576 mm^3^) exceeded threshold. ^b^ 91% probability of lying within the Harvard-Oxford amygdala. ^c^ 39% probability of lying within the Harvard-Oxford amygdala.

**Continued…**

**EXTENDED DATA FIG. 4-2**

**Descriptive statistics for clusters and local extrema showing greater activity during the anticipation of Uncertain Safety compared to Uncertain Threat (FDR *q*<.05, whole-brain corrected).**

|  | | **mm^3^** | ***t*** | ***x*** | ***y*** | ***z*** |
| --- | --- | --- | --- | --- | --- | --- |
|  | Cluster 1 | 51,072 |  |  |  |  |
|  | R Hippocampus |  | 3.80 | 24 | -20 | -20 |
|  | R Parahippocampal Gyrus, anterior division |  | 3.16 | 32 | -22 | -24 |
|  | R Parahippocampal Gyrus, posterior division |  | 3.18 | 28 | -28 | -22 |
|  | R Temporal Fusiform Cortex, posterior division |  | 5.15 | 28 | -36 | -16 |
|  | L Cingulate Gyrus, posterior division |  | 6.46 | -6 | -44 | 4 |
|  | R Cingulate Gyrus, posterior division |  | 8.04 | 6 | -46 | 4 |
|  | R Lingual Gyrus |  | 7.59 | 14 | -50 | 2 |
|  | R Precuneus Cortex |  | 10.10 | 4 | -60 | 20 |
|  | L Lingual Gyrus |  | 7.62 | -4 | -62 | 6 |
|  | L Precuneous Cortex |  | 7.64 | 0 | -62 | 34 |
|  | R Supracalcarine Cortex |  | 11.20 | 2 | -68 | 18 |
|  | R Cuneal Cortex |  | 10.66 | 6 | -70 | 22 |
|  | R Intracalcarine Cortex |  | 10.17 | 10 | -78 | 8 |
|  | L Intracalcarine Cortex |  | 11.11 | -14 | -84 | 4 |
|  | Cluster 2 | 41,664 |  |  |  |  |
|  | L Postcentral Gyrus |  | 9.59 | -52 | -10 | 26 |
|  | L Planum Polare |  | 3.42 | -50 | -10 | -2 |
|  | L Heschls Gyrus (*includes H1 and H2*) |  | 4.57 | -52 | -16 | 6 |
|  | L Central Opercular Cortex |  | 9.06 | -38 | -16 | 18 |
|  | L Parietal Operculum Cortex |  | 5.64 | -34 | -28 | 18 |
|  | R Precentral Gyrus |  | 10.36 | 2 | -30 | 60 |
|  | L Precentral Gyrus |  | 10.58 | -2 | -32 | 60 |
|  | R Postcentral Gyrus |  | 10.32 | 4 | -34 | 64 |
|  | Cluster 3 | 8,840 |  |  |  |  |
|  | R Insular Cortex |  | 7.67 | 36 | -10 | 16 |
|  | Cluster 4 | 8,096 |  |  |  |  |
|  | L Frontal Pole |  | 4.23 | -6 | 68 | 6 |
|  | R Inferior Rostral Sulcus |  | 5.62 | 4 | 54 | -6 |
|  | R Frontal Medial Cortex |  | 4.30 | 6 | 34 | -12 |
|  | R Subcallosal Cortex |  | 3.67 | 2 | 30 | -20 |
|  | L Subcallosal Cortex |  | 4.57 | -2 | 20 | -24 |
|  | Cluster 5 | 2,648 |  |  |  |  |
|  | R Lateral Occipital Cortex, superior division |  | 7.20 | 48 | -72 | 34 |
|  | Cluster 6 | 1,456 |  |  |  |  |
|  | L Lateral Occipital Cortex, superior division |  | 3.70 | -40 | -80 | 40 |
|  | Cluster 7 | 1,432 |  |  |  |  |
|  | L Parahippocampal Gyrus, posterior division |  | 5.64 | -26 | -38 | -16 |
|  | Cluster 8 | 920 |  |  |  |  |
|  | R Superior Frontal Gyrus |  | 4.80 | 22 | 32 | 48 |
|  | Cluster 9 | 392 |  |  |  |  |
|  | R Planum Temporale |  | 3.78 | 60 | -10 | 4 |
|  | Cluster 10 | 352 |  |  |  |  |
|  | L Middle Temporal Gyrus, posterior division |  | 3.97 | -60 | -12 | -14 |
|  | Cluster 11 | 288 |  |  |  |  |
|  | L Hippocampus |  | 3.91 | -18 | -12 | -22 |
|  | Cluster 12 | 264 |  |  |  |  |
|  | R Middle Temporal Gyrus, anterior division |  | 3.34 | 64 | -4 | -20 |
|  | Cluster 13 | 208 |  |  |  |  |
|  | L Thalamus |  | 3.12 | -18 | -26 | -2 |
|  | Cluster 14 | 192 |  |  |  |  |
|  | L Amygdala^a,b^ |  | 3.92 | -30 | 0 | -16 |
|  | Cluster 15 | 144 |  |  |  |  |
|  | L Frontal Medial Cortex |  | 3.39 | -2 | 44 | -26 |
|  | Cluster 16 | 96 |  |  |  |  |
|  | L Frontal Orbital Cortex |  | 3.57 | -34 | 36 | -10 |
|  | Cluster 17 | 96 |  |  |  |  |
|  | L Planum Temporale |  | 3.37 | -44 | -30 | 6 |
|  | Cluster 18 | 72 |  |  |  |  |
|  | R Amygdala^a,c^ |  | 3.61 | 28 | 2 | -16 |
|  | Cluster 19 | 64 |  |  |  |  |
|  | L Paracingulate Gyrus |  | 3.07 | -8 | 42 | -4 |
|  | Cluster 20 | 64 |  |  |  |  |
|  | L Middle Frontal Gyrus |  | 3.25 | -32 | 24 | 56 |
|  | Cluster 21 | 24 |  |  |  |  |
|  | L Juxtapositional Lobule Cortex |  | 2.99 | -10 | -14 | 46 |
|  | Cluster 22 | 8 |  |  |  |  |
|  | R Inferior Temporal Gyrus, posterior division |  | 2.81 | 54 | -32 | -30 |
|  | Cluster 23 | 8 |  |  |  |  |
|  | R Planum Polare |  | 2.87 | 52 | -4 | -6 |
|  | Cluster 24 | 8 |  |  |  |  |
|  | L Superior Temporal Gyrus, anterior division |  | 3.02 | -50 | -14 | -4 |
|  | Cluster 25 | 8 |  |  |  |  |
|  | R Thalamus |  | 2.81 | 16 | -28 | -2 |
|  | Cluster 26 | 8 |  |  |  |  |
|  | R Heschls Gyrus (*includes H1 and H2*) |  | 2.96 | 50 | -14 | 2 |

^a^ Within the Harvard-Oxford amygdala (*p*>.25), a total of 22 voxels (176 mm^3^) exceeded threshold. ^b^ 60% probability of lying within the Harvard-Oxford amygdala. ^c^ 61% probability of lying within the Harvard-Oxford amygdala.

**Continued…**

**EXTENDED DATA FIG. 4-3**

**Descriptive statistics for clusters and local extrema showing greater activity during the anticipation of Certain Threat compared to Certain Safety (FDR *q*<.05, whole-brain corrected).**

|  | | **mm^3^** | ***t*** | ***x*** | ***y*** | ***z*** |
| --- | --- | --- | --- | --- | --- | --- |
|  | Cluster 1 | 494,696 |  |  |  |  |
|  | L Frontal Pole |  | 7.08 | -38 | 54 | 14 |
|  | L Anterior Orbital Gyrus/Lateral Orbital Sulcus  (OP11) |  | 3.08 | -32 | 48 | -12 |
|  | L Middle Frontal Gyrus |  | 5.94 | -38 | 36 | 38 |
|  | R Frontal Pole |  | 5.03 | 32 | 36 | 28 |
|  | L Inferior Frontal Gyrus, pars triangularis |  | 5.64 | -54 | 34 | 4 |
|  | L Medial Orbital Gyrus |  | 4.07 | -18 | 34 | -16 |
|  | R Middle Frontal Gyrus |  | 4.43 | 34 | 34 | 34 |
|  | R Frontal Orbital Cortex |  | 3.76 | 40 | 32 | -20 |
|  | R Inferior Frontal Gyrus, pars triangularis |  | 3.79 | 44 | 32 | 2 |
|  | R Paracingulate Gyrus |  | 4.06 | 14 | 30 | 26 |
|  | L Frontal Orbital Cortex |  | 4.96 | -36 | 28 | -4 |
|  | L Cingulate Gyrus, anterior division |  | 6.64 | -6 | 20 | 34 |
|  | L Frontal Operculum Cortex |  | 4.59 | -38 | 18 | 10 |
|  | R Frontal Operculum Cortex |  | 6.47 | 50 | 18 | 0 |
|  | R Cingulate Gyrus, anterior division |  | 7.07 | 6 | 16 | 36 |
|  | R Inferior Frontal Gyrus, pars opercularis |  | 6.53 | 52 | 16 | -2 |
|  | L Inferior Frontal Gyrus, pars opercularis |  | 3.09 | -52 | 14 | 14 |
|  | L Temporal Pole |  | 7.83 | -52 | 12 | -6 |
|  | L Paracingulate Gyrus |  | 6.18 | -2 | 12 | 46 |
|  | L Caudate |  | 5.84 | -16 | 10 | 16 |
|  | L Central Opercular Cortex |  | 2.68 | -32 | 8 | 16 |
|  | R Accumbens |  | 3.63 | 12 | 8 | -6 |
|  | R Caudate |  | 6.59 | 18 | 8 | 20 |
|  | R Putamen |  | 6.29 | 22 | 8 | -8 |
|  | L Putamen |  | 6.51 | -18 | 6 | -10 |
|  | R Temporal Pole |  | 5.03 | 50 | 6 | -40 |
|  | L Bed Nucleus of the Stria Terminalis |  | 3.75 | -8 | 2 | 2 |
|  | R Bed Nucleus of the Stria Terminalis |  | 3.75 | 10 | 2 | 2 |
|  | L Superior Frontal Gyrus |  | 6.47 | -18 | 0 | 70 |
|  | R Juxtapositional Lobule Cortex |  | 7.02 | 4 | 0 | 68 |
|  | R Superior Frontal Gyrus |  | 6.44 | 16 | 0 | 68 |
|  | L Superior Temporal Gyrus, anterior division |  | 4.28 | -64 | -2 | 0 |
|  | L Middle Temporal Gyrus, anterior division |  | 4.95 | -56 | -2 | -36 |
|  | L Parahippocampal Gyrus, anterior division / Amygdala |  | 3.66 | -22 | -2 | -28 |
|  | L Pallidum |  | 4.62 | -16 | -2 | 4 |
|  | L Juxtapositional Lobule Cortex |  | 6.76 | -2 | -2 | 66 |
|  | L Precentral Gyrus |  | 6.86 | -30 | -4 | 64 |
|  | L Temporal Fusiform Cortex, anterior division |  | 3.41 | -30 | -6 | -34 |
|  | R Inferior Temporal Gyrus, anterior division |  | 3.16 | 44 | -6 | -36 |
|  | R Pallidum |  | 5.13 | 18 | -8 | -2 |
|  | L Amygdala (*in the region of the Cortical nucleus*  *and Amygdala-Hippocampal Transition Area*)^a,b^ |  | 2.86 | -32 | 0 | -22 |
|  | L Inferior Temporal Gyrus, anterior division |  | 2.88 | -44 | -10 | -34 |
|  | R Precentral Gyrus |  | 7.20 | 30 | -10 | 66 |
|  | R Amygdala/Hippocampus^a,c^ |  | 5.33 | 18 | -12 | -14 |
|  | L Inferior Temporal Gyrus, posterior division |  | 4.77 | -56 | -14 | -30 |
|  | R Hippocampus |  | 4.97 | 30 | -16 | -12 |
|  | L Superior Temporal Gyrus, posterior division |  | 3.11 | -64 | -18 | 2 |
|  | L Hippocampus |  | 4.61 | -20 | -22 | -16 |
|  | R Cingulate Gyrus, posterior division |  | 4.34 | 6 | -24 | 42 |
|  | R Inferior Temporal Gyrus, posterior division |  | 4.14 | 52 | -24 | -22 |
|  | L Postcentral Gyrus |  | 2.67 | -46 | -26 | 46 |
|  | L Middle Temporal Gyrus, posterior division |  | 4.47 | -66 | -28 | -6 |
|  | L Cingulate Gyrus, posterior division |  | 4.59 | -12 | -30 | 36 |
|  | L Supramarginal Gyrus, anterior division |  | 3.56 | -54 | -32 | 34 |
|  | L Parahippocampal Gyrus, posterior division |  | 5.35 | -14 | -32 | -12 |
|  | R Middle Temporal Gyrus, posterior division |  | 5.27 | 48 | -32 | -6 |
|  | R Supramarginal Gyrus, anterior division |  | 4.68 | 50 | -32 | 56 |
|  | R Superior Temporal Gyrus, posterior division |  | 3.36 | 58 | -32 | 10 |
|  | L Brainstem/Periaqueductal Gray |  | 2.66 | -2 | -34 | -14 |
|  | L Temporal Fusiform Cortex, posterior division |  | 4.16 | -30 | -34 | -28 |
|  | R Thalamus |  | 5.65 | 10 | -34 | 10 |
|  | R Temporal Fusiform Cortex, posterior division |  | 2.94 | 40 | -34 | -18 |
|  | R Parietal Operculum Cortex |  | 3.20 | 56 | -34 | 26 |
|  | L Thalamus |  | 5.55 | -22 | -36 | -2 |
|  | R Planum Temporale |  | 3.30 | 50 | -36 | 16 |
|  | L Planum Temporale |  | 3.29 | -64 | -38 | 18 |
|  | R Postcentral Gyrus |  | 5.85 | 38 | -38 | 66 |
|  | R Middle Temporal Gyrus, temporooccipital part |  | 6.14 | 50 | -38 | 2 |
|  | R Superior Parietal Lobule |  | 5.91 | 34 | -44 | 62 |
|  | R Supramarginal Gyrus, posterior division |  | 7.25 | 56 | -44 | 34 |
|  | L Supramarginal Gyrus, posterior division |  | 6.17 | -62 | -46 | 32 |
|  | L Precuneus Cortex |  | 3.94 | -8 | -48 | 48 |
|  | R Temporal Occipital Fusiform Cortex |  | 2.79 | 42 | -52 | -16 |
|  | L Angular Gyrus |  | 7.51 | -56 | -54 | 44 |
|  | L Inferior Temporal Gyrus, temporooccipital part |  | 5.05 | -56 | -54 | -20 |
|  | L Superior Parietal Lobule |  | 6.90 | -40 | -54 | 62 |
|  | L Middle Temporal Gyrus, temporooccipital part |  | 5.28 | -48 | -56 | 6 |
|  | R Angular Gyrus |  | 3.76 | 38 | -56 | 36 |
|  | R Inferior Temporal Gyrus, temporooccipital part |  | 4.18 | 54 | -58 | -14 |
|  | L Lateral Occipital Cortex, superior division |  | 6.36 | -30 | -60 | 62 |
|  | R Precuneus Cortex |  | 5.93 | 8 | -62 | 56 |
|  | R Lateral Occipital Cortex, superior division |  | 6.59 | 14 | -68 | 60 |
|  | R Lateral Occipital Cortex, inferior division |  | 4.48 | 48 | -76 | -12 |
|  | L Cuneal Cortex |  | 3.81 | -18 | -78 | 30 |
|  | R Lingual Gyrus |  | 5.82 | 4 | -78 | -16 |
|  | L Occipital Fusiform Gyrus |  | 5.76 | -8 | -84 | -20 |
|  | R Occipital Pole |  | 4.36 | 2 | -94 | -18 |
|  | Cluster 2 | 600 |  |  |  |  |
|  | L Lateral Occipital Cortex, inferior division |  | 3.64 | -48 | -80 | -8 |
|  | Cluster 3 | 280 |  |  |  |  |
|  | R Planum Polare |  | 3.20 | 42 | -8 | -14 |
|  | Cluster 4 | 160 |  |  |  |  |
|  | R Parahippocampal Gyrus, anterior division |  | 2.94 | 20 | -14 | -32 |
|  | Cluster 5 | 160 |  |  |  |  |
|  | R Subcallosal Cortex |  | 2.86 | 10 | 16 | -22 |
|  | Cluster 6 | 48 |  |  |  |  |
|  | R Middle Temporal Gyrus, anterior division |  | 3.12 | 50 | -2 | -26 |
|  | Cluster 7 | 48 |  |  |  |  |
|  | L Lingual Gyrus |  | 2.46 | -20 | -52 | 0 |
|  | Cluster 8 | 40 |  |  |  |  |
|  | L Frontal Medial Cortex |  | 2.67 | -4 | 52 | -28 |
|  | Cluster 9 | 40 |  |  |  |  |
|  | R Central Opercular Cortex |  | 2.61 | 38 | -10 | 22 |
|  | Cluster 10 | 32 |  |  |  |  |
|  | L Planum Polare |  | 2.51 | -40 | -4 | -16 |
|  | Cluster 11 | 16 |  |  |  |  |
|  | L Parietal Operculum Cortex |  | 2.80 | -54 | -24 | 20 |
|  | Cluster 12 | 16 |  |  |  |  |
|  | R Cuneal Cortex |  | 2.34 | 16 | -80 | 28 |

^a^ Within the Harvard-Oxford amygdala (*p*>.25), a total of 148 voxels (1,184 mm^3^) exceeded threshold. ^b^ 41% probability of lying within the Harvard-Oxford amygdala. ^c^ 57% probability of lying within the Harvard-Oxford amygdala.

**Continued…**

**EXTENDED DATA FIG. 4-4**

**Descriptive statistics for clusters and local extrema showing greater activity during the anticipation of Certain Safety compared to Certain Threat (FDR *q*<.05, whole-brain corrected).**

|  | | **mm^3^** | ***t*** | ***x*** | ***y*** | ***z*** |
| --- | --- | --- | --- | --- | --- | --- |
|  | Cluster 1 | 63,216 |  |  |  |  |
|  | R Parahippocampal Gyrus, posterior division |  | 4.98 | 22 | -36 | -12 |
|  | R Lingual Gyrus |  | 6.70 | 20 | -46 | -10 |
|  | R Cingulate Gyrus, posterior division |  | 4.44 | 2 | -48 | 20 |
|  | L Cingulate Gyrus, posterior division |  | 3.07 | -6 | -50 | 10 |
|  | L Precuneus Cortex |  | 3.39 | -2 | -54 | 6 |
|  | R Temporal Occipital Fusiform Cortex |  | 5.42 | 34 | -56 | -14 |
|  | L Supracalcarine Cortex |  | 10.57 | 0 | -68 | 14 |
|  | R Cuneal Cortex |  | 9.02 | 8 | -68 | 20 |
|  | R Occipital Fusiform Gyrus |  | 6.75 | 30 | -74 | -8 |
|  | R Intracalcarine Cortex |  | 11.55 | 12 | -80 | 4 |
|  | L Occipital Fusiform Gyrus |  | 5.24 | -26 | -86 | -16 |
|  | L Cuneal Cortex |  | 4.00 | 0 | -86 | 28 |
|  | L Intracalcarine Cortex |  | 11.84 | -12 | -88 | 4 |
|  | R Occipital Pole |  | 11.16 | 6 | -90 | 6 |
|  | L Occipital Pole |  | 9.10 | -8 | -100 | -2 |
|  | Cluster 2 | 3,864 |  |  |  |  |
|  | R Precentral Gyrus |  | 5.25 | 4 | -28 | 54 |
|  | L Precentral Gyrus |  | 5.27 | -4 | -30 | 58 |
|  | Cluster 3 | 2,688 |  |  |  |  |
|  | L Postcentral Gyrus |  | 5.39 | -52 | -10 | 26 |
|  | Cluster 5 | 2,240 |  |  |  |  |
|  | L Insular Cortex |  | 4.29 | -36 | -12 | 10 |
|  | L Central Opercular Cortex |  | 6.39 | -38 | -18 | 18 |
|  | L Heschl’s Gyrus (includes H1 and H2) |  | 3.44 | -36 | -24 | 8 |
|  | L Parietal Operculum Cortex |  | 5.97 | -34 | -28 | 18 |
|  | Cluster 6 | 1,096 |  |  |  |  |
|  | R Insular Cortex |  | 5.97 | 34 | -26 | 16 |
|  | Cluster 8 | 864 |  |  |  |  |
|  | L Lingual Gyrus |  | 4.28 | -18 | -50 | -10 |
|  | Cluster 12 | 328 |  |  |  |  |
|  | R Central Opercular Cortex |  | 3.35 | 50 | -10 | 14 |
|  | R Heschls Gyrus (includes H1 and H2) |  | 3.98 | 46 | -14 | 6 |
|  | Cluster 13 | 328 |  |  |  |  |
|  | L Juxtapositional Lobule Cortex |  | 4.33 | -10 | -10 | 42 |
|  | Cluster 14 | 256 |  |  |  |  |
|  | R Cingulate Gyrus, anterior division |  | 3.87 | 2 | 34 | -4 |
|  | Cluster 16 | 232 |  |  |  |  |
|  | R Postcentral Gyrus |  | 4.58 | 54 | -14 | 56 |
|  | Cluster 19 | 144 |  |  |  |  |
|  | L Superior Temporal Gyrus, anterior division |  | 3.75 | -50 | -4 | -16 |
|  | Cluster 21 | 56 |  |  |  |  |
|  | R Subcallosal Cortex |  | 3.35 | 0 | 24 | -20 |
|  | Cluster 22 | 56 |  |  |  |  |
|  | R Superior Frontal Gyrus |  | 3.42 | 24 | 18 | 46 |
|  | Cluster 23 | 48 |  |  |  |  |
|  | L Frontal Orbital Cortex |  | 3.47 | -28 | 10 | -20 |
|  | Cluster 24 | 40 |  |  |  |  |
|  | L Parahippocampal Gyrus, posterior division |  | 3.22 | -32 | -34 | -14 |
|  | Cluster 26 | 24 |  |  |  |  |
|  | R Frontal Pole |  | 3.17 | 8 | 62 | 0 |
|  | Cluster 27 | 24 |  |  |  |  |
|  | R Precuneus Cortex |  | 3.29 | 24 | -58 | 10 |
|  | Cluster 29 | 16 |  |  |  |  |
|  | L Temporal Occipital Fusiform Cortex |  | 3.16 | -34 | -48 | -16 |

**Continued…
EXTENDED DATA FIG. 4-5**

**Descriptive statistics for clusters and local extrema identified by the minimum conjunction (logical ‘AND’) of the Uncertain Threat greater than Uncertain Safety and Certain Threat greater than Certain Safety contrasts (FDR *q*<.05, whole-brain corrected).**

|  | | **mm^3^** | ***t_Minimum_*** | ***x*** | ***y*** | ***z*** |
| --- | --- | --- | --- | --- | --- | --- |
|  | Cluster 1 | 259,760 |  |  |  |  |
|  | L Frontal Pole |  | 6.83 | -30 | 54 | 28 |
|  | L Anterior Orbital Gyrus/Lateral Orbital Sulcus  (OP11) |  | 2.89 | -30 | 48 | -12 |
|  | L Brainstem/Periaqueductal Gray |  | 2.66 | -2 | 34 | -14 |
|  | R Inferior Frontal Gyrus, pars triangularis |  | 2.99 | 48 | 30 | 4 |
|  | L Frontal Orbital Cortex |  | 4.96 | -36 | 28 | -4 |
|  | R Frontal Orbital Cortex |  | 3.64 | 38 | 24 | -12 |
|  | L Insular Cortex |  | 5.06 | -28 | 22 | 10 |
|  | L Cingulate Gyrus, anterior division |  | 6.64 | -6 | 20 | 34 |
|  | L Frontal Operculum Cortex |  | 4.59 | -38 | 18 | 10 |
|  | R Frontal Operculum Cortex |  | 6.47 | 50 | 18 | 0 |
|  | R Cingulate Gyrus, anterior division |  | 7.07 | 6 | 16 | 36 |
|  | R Inferior Frontal Gyrus, pars opercularis |  | 6.53 | 52 | 16 | -2 |
|  | L Inferior Frontal Gyrus, pars opercularis |  | 3.09 | -52 | 14 | 14 |
|  | L Temporal Pole |  | 7.78 | -52 | 12 | -4 |
|  | L Caudate |  | 5.12 | -18 | 12 | 14 |
|  | L Paracingulate Gyrus |  | 6.18 | -2 | 12 | 46 |
|  | L Putamen |  | 5.86 | -24 | 10 | -4 |
|  | R Accumbens |  | 2.80 | 10 | 8 | -4 |
|  | R Caudate |  | 6.11 | 18 | 8 | 18 |
|  | R Putamen |  | 6.18 | 22 | 8 | -6 |
|  | L Accumbens |  | 2.62 | -8 | 6 | -4 |
|  | L Bed Nucleus of the Stria Terminalis |  | 3.70 | -8 | 4 | 2 |
|  | L Superior Frontal Gyrus |  | 6.47 | -18 | 0 | 70 |
|  | R Juxtapositional Lobule Cortex |  | 7.02 | 4 | 0 | 68 |
|  | R Superior Frontal Gyrus |  | 6.44 | 16 | 0 | 68 |
|  | L Pallidum |  | 4.50 | -16 | -2 | 2 |
|  | L Juxtapositional Lobule Cortex |  | 6.49 | 0 | -2 | 64 |
|  | R Amygdala (*in the region of the Central and Medial*  *nuclei*)^a,b^ |  | 2.36 | 20 | -4 | -10 |
|  | L Precentral Gyrus |  | 6.38 | -34 | -6 | 62 |
|  | R Pallidum |  | 4.71 | 18 | -6 | -2 |
|  | R Amygdala/Lentiform Nucleus^a^ |  | 2.51 | 30 | -8 | -16 |
|  | L Thalamus |  | 5.12 | 0 | -10 | 10 |
|  | R Precentral Gyrus |  | 7.20 | 30 | -10 | 66 |
|  | R Thalamus |  | 4.36 | 6 | -24 | 14 |
|  | L Temporal Fusiform Cortex, posterior division |  | 2.59 | -36 | -32 | -24 |
|  | L Parahippocampal Gyrus, posterior division |  | 4.57 | -12 | -32 | -12 |
|  | L Inferior Temporal Gyrus, temporooccipital part |  | 4.46 | -48 | -46 | -28 |
|  | R Temporal Occipital Fusiform Cortex |  | 2.37 | 42 | -52 | -16 |
|  | R Lingual Gyrus |  | 5.82 | 4 | -78 | -16 |
|  | Cluster 2 | 79,048 |  |  |  |  |
|  | L Middle Frontal Gyrus |  | 5.94 | -38 | 36 | 38 |
|  | L Cingulate Gyrus, posterior division |  | 3.64 | -4 | -24 | 42 |
|  | R Cingulate Gyrus, posterior division |  | 4.34 | 6 | -24 | 42 |
|  | R Supramarginal Gyrus, anterior division |  | 4.16 | 50 | -30 | 42 |
|  | R Middle Temporal Gyrus, posterior division |  | 5.27 | 48 | -32 | -6 |
|  | R Planum Temporale |  | 2.64 | 54 | -34 | 20 |
|  | R Parietal Operculum Cortex |  | 3.20 | 56 | -34 | 26 |
|  | L Postcentral Gyrus |  | 3.59 | -46 | -36 | 50 |
|  | R Postcentral Gyrus |  | 4.55 | 36 | -36 | 50 |
|  | L Planum Temporale |  | 3.29 | -64 | -38 | 18 |
|  | R Middle Temporal Gyrus, temporooccipital part |  | 6.14 | 48 | -38 | 2 |
|  | L Supramarginal Gyrus, anterior division |  | 4.73 | -54 | -40 | 38 |
|  | R Supramarginal Gyrus, posterior division |  | 7.25 | 56 | -44 | 34 |
|  | L Supramarginal Gyrus, posterior division |  | 6.17 | -62 | -46 | 32 |
|  | R Superior Parietal Lobule |  | 5.67 | 32 | -46 | 66 |
|  | R Angular Gyrus |  | 2.84 | 58 | -46 | 14 |
|  | L Superior Parietal Lobule |  | 5.36 | -34 | -50 | 64 |
|  | L Angular Gyrus |  | 7.06 | -56 | -52 | 42 |
|  | R Precuneus Cortex |  | 4.54 | 10 | -56 | 54 |
|  | R Lateral Occipital Cortex, superior division |  | 3.83 | 16 | -58 | 66 |
|  | L Precuneus Cortex |  | 3.92 | -10 | -72 | 34 |
|  | L Lateral Occipital Cortex, superior division |  | 4.43 | -12 | -76 | 46 |
|  | L Cuneal Cortex |  | 3.57 | -16 | -80 | 32 |
|  | Cluster 3 | 10,152 |  |  |  |  |
|  | R Frontal Pole |  | 5.03 | 32 | 36 | 28 |
|  | R Middle Frontal Gyrus |  | 4.43 | 34 | 34 | 34 |
|  | Cluster 4 | 3,848 |  |  |  |  |
|  | L Middle Temporal Gyrus, temporooccipital part |  | 3.75 | -60 | -58 | 6 |
|  | L Lateral Occipital Cortex, inferior division |  | 3.15 | -52 | -64 | 12 |
|  | Cluster 5 | 3,296 |  |  |  |  |
|  | R Temporal Pole |  | 4.55 | 48 | 6 | -42 |
|  | R Middle Temporal Gyrus, anterior division |  | 2.37 | 56 | -2 | -32 |
|  | R Temporal Fusiform Cortex, posterior division |  | 2.48 | 44 | -10 | -34 |
|  | R Inferior Temporal Gyrus, posterior division |  | 3.26 | 58 | -20 | -32 |
|  | Cluster 6 | 2,152 |  |  |  |  |
|  | L Inferior Temporal Gyrus, anterior division |  | 3.58 | -52 | -2 | -36 |
|  | L Middle Temporal Gyrus anterior division |  | 3.64 | -54 | -4 | -34 |
|  | L Inferior Temporal Gyrus, posterior division |  | 3.19 | -56 | -18 | -30 |
|  | L Middle Temporal Gyrus, posterior division |  | 3.79 | -58 | -30 | -6 |
|  | Cluster 7 | 1,248 |  |  |  |  |
|  | R Cuneal Cortex |  | 3.81 | 16 | -78 | 36 |
|  | Cluster 8 | 736 |  |  |  |  |
|  | R Right Hippocampus |  | 3.52 | 30 | -38 | -2 |
|  | Cluster 9 | 576 |  |  |  |  |
|  | R Occipital Pole |  | 3.52 | 32 | -88 | 12 |
|  | Cluster 10 | 280 |  |  |  |  |
|  | L Temporal Fusiform Cortex, anterior division |  | 3.05 | -32 | -4 | -34 |
|  | L Parahippocampal Gyrus, anterior division |  | 2.66 | -26 | -6 | -30 |
|  | Cluster 11 | 264 |  |  |  |  |
|  | R Insular Cortex |  | 3.00 | 42 | -6 | -8 |
|  | R Planum Polare |  | 3.20 | 42 | -8 | -14 |
|  | Cluster 12 | 232 |  |  |  |  |
|  | R Lateral Occipital Cortex, inferior division |  | 3.26 | 46 | -66 | 12 |
|  | Cluster 13 | 224 |  |  |  |  |
|  | L Hippocampus |  | 3.49 | -34 | -24 | -10 |
|  | Cluster 14 | 112 |  |  |  |  |
|  | R Parahippocampal Gyrus, anterior division |  | 2.94 | 20 | -14 | -32 |
|  | Cluster 15 | 56 |  |  |  |  |
|  | R Inferior Temporal Gyrus, temporooccipital part |  | 2.29 | 58 | -52 | -16 |
|  | Cluster 16 | 32 |  |  |  |  |
|  | R Inferior Temporal Gyrus, anterior division |  | 2.40 | 40 | -4 | -42 |
|  | Cluster 17 | 32 |  |  |  |  |
|  | L Lingual Gyrus |  | 2.27 | -20 | -52 | -2 |
|  | Cluster 18 | 32 |  |  |  |  |
|  | R Paracingulate Gyrus |  | 2.42 | 10 | 46 | 4 |
|  | Cluster 19 | 16 |  |  |  |  |
|  | L Planum Polare |  | 2.51 | -40 | -4 | -16 |
|  | Cluster 20 | 16 |  |  |  |  |
|  | L Central Opercular Cortex |  | 2.56 | -32 | 8 | 16 |
|  | Cluster 21 | 16 |  |  |  |  |
|  | L Parietal Operculum Cortex |  | 2.80 | -54 | -24 | 20 |
|  | Cluster 22 | 8 |  |  |  |  |
|  | R Subcallosal Cortex |  | 2.12 | 10 | 18 | -22 |

^a^Within the Harvard-Oxford amygdala (*p*>.25), a total of 13 voxels (104 mm^3^) exceeded threshold. ^b^ 40% probability of lying within the Harvard-Oxford amygdala.

**Continued…**

**EXTENDED DATA FIG. 7-1**

**Descriptive statistics for clusters and local extrema showing greater activity during the anticipation of Uncertain compared to Certain Threat (FDR *q*<.05, whole-brain corrected).**

|  | | **mm^3^** | ***t*** | ***x*** | ***y*** | ***z*** |
| --- | --- | --- | --- | --- | --- | --- |
|  | Cluster 1 | 116,856 |  |  |  |  |
|  | R Paracingulate Gyrus |  | 5.10 | 2 | 40 | 24 |
|  | R Frontal Pole |  | 4.79 | 52 | 40 | -6 |
|  | L Paracingulate Gyrus |  | 5.61 | -6 | 34 | 26 |
|  | R Inferior Frontal Gyrus, pars triangularis |  | 2.72 | 46 | 32 | 6 |
|  | R Frontal Orbital Cortex |  | 9.73 | 38 | 30 | 0 |
|  | L Cingulate Gyrus, anterior division |  | 6.49 | -4 | 24 | 32 |
|  | R Frontal Operculum Cortex |  | 9.35 | 36 | 24 | 4 |
|  | R Superior Frontal Gyrus |  | 6.87 | 8 | 22 | 60 |
|  | R Inferior Frontal Gyrus, pars opercularis |  | 8.77 | 52 | 20 | 4 |
|  | R Middle Frontal Gyrus |  | 4.31 | 46 | 18 | 42 |
|  | L Superior Frontal Gyrus |  | 6.10 | -14 | 12 | 62 |
|  | R Insular Cortex |  | 6.25 | 34 | 6 | 6 |
|  | R Cingulate Gyrus, anterior division |  | 8.53 | 4 | 2 | 40 |
|  | R Central Opercular Cortex |  | 8.02 | 52 | 2 | 6 |
|  | R Precentral Gyrus |  | 8.22 | 46 | 0 | 50 |
|  | R Putamen |  | 5.27 | 28 | -2 | 8 |
|  | R Pallidum |  | 4.09 | 24 | -6 | -6 |
|  | R Heschls Gyrus |  | 5.51 | 46 | -12 | 6 |
|  | R Cingulate Gyrus, posterior division |  | 5.95 | 2 | -18 | 40 |
|  | L Cingulate Gyrus, posterior division |  | 4.55 | -4 | -24 | 26 |
|  | R Parietal Operculum Cortex |  | 7.26 | 54 | -26 | 24 |
|  | R Supramarginal Gyrus, posterior division |  | 5.62 | 60 | -42 | 30 |
|  | R Angular Gyrus |  | 5.58 | 58 | -50 | 36 |
|  | Cluster 2 | 68,280 |  |  |  |  |
|  | L Temporal Occipital Fusiform Cortex |  | 3.64 | -34 | -48 | -16 |
|  | R Temporal Occipital Fusiform Cortex |  | 7.74 | 34 | -52 | -16 |
|  | R Occipital Fusiform Gyrus |  | 9.60 | 36 | -64 | -14 |
|  | R Lateral Occipital Cortex, inferior division |  | 12.89 | 38 | -82 | -10 |
|  | R Lingual Gyrus |  | 4.77 | 10 | -86 | -10 |
|  | L Occipital Fusiform Gyrus |  | 6.78 | -22 | -88 | -8 |
|  | L Lateral Occipital Cortex, inferior division |  | 5.14 | -38 | -90 | -12 |
|  | L Occipital Pole |  | 7.90 | -14 | -98 | -4 |
|  | R Occipital Pole |  | 10.04 | 18 | -98 | 2 |
|  | Cluster 3 | 21,816 |  |  |  |  |
|  | L Frontal Orbital Cortex |  | 10.39 | -32 | 26 | 0 |
|  | L Inferior Frontal Gyrus, pars opercularis |  | 6.85 | -50 | 20 | 4 |
|  | L Precentral Gyrus |  | 3.96 | -54 | 2 | 12 |
|  | L Central Opercular Cortex |  | 5.54 | -48 | -2 | 6 |
|  | L Planum Polare |  | 2.95 | -42 | -2 | -14 |
|  | L Putamen |  | 5.36 | -34 | -2 | 0 |
|  | Cluster 4 | 8,640 |  |  |  |  |
|  | R Thalamus |  | 5.17 | 12 | -10 | 4 |
|  | Cluster 5 | 6,784 |  |  |  |  |
|  | L Supramarginal Gyrus, anterior division |  | 3.99 | -64 | -26 | 22 |
|  | L Parietal Operculum Cortex |  | 4.67 | -56 | -32 | 24 |
|  | L Supramarginal Gyrus, posterior division |  | 4.69 | -62 | -52 | 38 |
|  | L Angular Gyrus |  | 3.97 | -54 | -56 | 36 |
|  | Cluster 6 | 4,720 |  |  |  |  |
|  | R Superior Parietal Lobule |  | 7.64 | 20 | -48 | 66 |
|  | Cluster 7 | 3,336 |  |  |  |  |
|  | L Frontal Pole |  | 4.70 | -36 | 40 | 30 |
|  | Cluster 8 | 1,616 |  |  |  |  |
|  | L Middle Frontal Gyrus |  | 3.55 | -42 | 8 | 48 |
|  | Cluster 9 | 672 |  |  |  |  |
|  | R Temporal Pole |  | 4.01 | 42 | 6 | -34 |
|  | R Middle Temporal Gyrus, anterior division |  | 2.98 | 46 | 2 | -28 |
|  | Cluster 10 | 664 |  |  |  |  |
|  | L Superior Parietal Lobule |  | 3.79 | -20 | -52 | 64 |
|  | L Precuneus Cortex |  | 2.76 | -10 | -52 | 58 |
|  | Cluster 11 | 472 |  |  |  |  |
|  | R Middle Temporal Gyrus, temporooccipital part |  | 4.03 | 56 | -44 | 4 |
|  | Cluster 12 | 344 |  |  |  |  |
|  | L Thalamus |  | 3.50 | -10 | -12 | 10 |
|  | Cluster 13 | 224 |  |  |  |  |
|  | L Insular Cortex |  | 3.86 | -38 | -22 | -2 |
|  | Cluster 14 | 192 |  |  |  |  |
|  | L Temporal Pole |  | 2.96 | -40 | 2 | -40 |
|  | Cluster 15 | 152 |  |  |  |  |
|  | R Planum Polare |  | 3.66 | 42 | 0 | -20 |
|  | Cluster 16 | 144 |  |  |  |  |
|  | R Superior Temporal Gyrus, posterior division |  | 2.88 | 50 | -20 | -4 |
|  | Cluster 17 | 112 |  |  |  |  |
|  | R Inferior Temporal Gyrus, posterior division |  | 3.19 | 52 | -36 | -22 |
|  | Cluster 18 | 104 |  |  |  |  |
|  | R Inferior Temporal Gyrus, temporooccipital part |  | 3.67 | 64 | -40 | -22 |
|  | Cluster 19 | 80 |  |  |  |  |
|  | R Middle Temporal Gyrus, posterior division |  | 3.05 | 60 | -28 | -6 |
|  | Cluster 20 | 48 |  |  |  |  |
|  | L Pallidum |  | 3.04 | -14 | -4 | -6 |
|  | Cluster 21 | 40 |  |  |  |  |
|  | R Hippocampus |  | 3.87 | 28 | -42 | 2 |
|  | Cluster 22 | 32 |  |  |  |  |
|  | L Inferior Temporal Gyrus, anterior division |  | 2.83 | -46 | 0 | -36 |
|  | Cluster 23 | 24 |  |  |  |  |
|  | L Temporal Fusiform Cortex, posterior division |  | 3.31 | -34 | -16 | -40 |

**Continued…**

**EXTENDED DATA FIG. 7-2**

**Descriptive statistics for clusters and local extrema showing greater activity during the anticipation of Certain compared to Uncertain Threat (FDR *q*<.05, whole-brain corrected).**

|  | | **mm^3^** | ***t*** | ***x*** | ***y*** | ***z*** |
| --- | --- | --- | --- | --- | --- | --- |
|  | Cluster 1 | 202,448 |  |  |  |  |
|  | L Caudate |  | 6.56 | -6 | 16 | 2 |
|  | L Putamen |  | 4.84 | -14 | 12 | -6 |
|  | R Putamen |  | 5.49 | 18 | 12 | -6 |
|  | L Subcallosal Cortex |  | 6.40 | -2 | 10 | -2 |
|  | R Caudate |  | 5.52 | 8 | 8 | 0 |
|  | R Bed Nucleus of the Stria Terminalis |  | 4.17 | 8 | 4 | 0 |
|  | L Temporal Pole |  | 4.44 | -34 | 4 | -22 |
|  | L Bed Nucleus of the Stria Terminalis |  | 4.39 | -4 | 4 | -2 |
|  | R Amygdala^a,b^ |  | 4.06 | 30 | 2 | -16 |
|  | L Amygdala^a,c^ |  | 3.80 | -32 | -2 | -16 |
|  | L Amygdala (*in the region of the Cortical nucleus and the Amygdala-Hippocampal Transition Area, medial to the Central nucleus)*^a,d^ |  | 4.83 | -18 | -10 | -14 |
|  | L Central Opercular Cortex |  | 7.21 | -36 | -12 | 16 |
|  | L Parahippocampal Gyrus, anterior division |  | 3.42 | -22 | -18 | -28 |
|  | L Hippocampus |  | 6.17 | -22 | -20 | -16 |
|  | R Hippocampus |  | 6.29 | 24 | -20 | -16 |
|  | R Parahippocampal Gyrus, anterior division |  | 3.55 | 24 | -20 | -28 |
|  | L Precentral Gyrus |  | 5.89 | -10 | -28 | 74 |
|  | R Precentral Gyrus |  | 6.74 | 4 | -30 | 66 |
|  | L Parahipppocampal Gyrus, posterior division |  | 4.64 | -26 | -32 | -20 |
|  | L Thalamus |  | 5.37 | -8 | -32 | 6 |
|  | R Thalamus |  | 6.40 | 10 | -34 | 8 |
|  | R Temporal Fusiform Cortex, posterior division |  | 4.10 | 24 | -36 | -24 |
|  | R Parahippocampal Gyrus, posterior division |  | 5.79 | 36 | -36 | -10 |
|  | L Temporal Fusiform Cortex, posterior division |  | 6.15 | -26 | -40 | -16 |
|  | L Postcentral Gyrus |  | 8.51 | -8 | -40 | 74 |
|  | L Cingulate Gyrus, posterior division |  | 7.47 | -6 | -44 | 4 |
|  | R Cingulate Gyrus, posterior division |  | 7.48 | 8 | -50 | 6 |
|  | L Lingual Gyrus |  | 6.24 | -16 | -54 | 2 |
|  | L Middle Temporal Gyrus, temporooccipital part |  | 3.66 | -54 | -56 | -8 |
|  | L Superior Parietal Lobule |  | 4.74 | -32 | -56 | 56 |
|  | R Lingual Gyrus |  | 9.19 | 10 | -58 | 6 |
|  | R Angular Gyrus |  | 3.41 | 48 | -58 | 16 |
|  | R Intracalcarine Cortex |  | 9.23 | 8 | -60 | 8 |
|  | R Precuneus Cortex |  | 7.43 | 16 | -60 | 18 |
|  | L Angular Gyrus |  | 3.61 | -42 | -62 | 24 |
|  | L Precuneus Cortex |  | 7.33 | -2 | -62 | 36 |
|  | L Lateral Occipital Cortex, inferior division |  | 4.24 | -44 | -64 | 0 |
|  | R Cuneal Cortex |  | 7.50 | 20 | -66 | 20 |
|  | L Cuneal Cortex |  | 7.90 | -10 | -70 | 18 |
|  | L Intracalcarine Cortex |  | 4.84 | 0 | -76 | 10 |
|  | R Lateral Occipital Cortex, superior division |  | 7.34 | 42 | -82 | 26 |
|  | L Lateral Occipital Cortex, superior division |  | 6.36 | -30 | -88 | 34 |
|  | R Occipital Pole |  | 3.42 | 8 | -90 | 26 |
|  | Cluster 2 | 14,480 |  |  |  |  |
|  | L Frontal Pole |  | 5.66 | -2 | 56 | -6 |
|  | R Frontal Pole |  | 5.56 | 4 | 54 | -6 |
|  | L Paracingulate Gyrus |  | 3.11 | -12 | 50 | -2 |
|  | R Frontal Medial Cortex |  | 4.87 | 0 | 48 | -14 |
|  | R Paracingulate Gyrus |  | 3.47 | 14 | 42 | -4 |
|  | L Frontal Medial Cortex |  | 3.27 | -4 | 36 | -16 |
|  | Cluster 3 | 9,256 |  |  |  |  |
|  | R Central Opercular Cortex |  | 4.01 | 44 | -8 | 18 |
|  | R Insular Cortex |  | 5.23 | 36 | -10 | 16 |
|  | R Postcentral Gyrus |  | 6.29 | 64 | -10 | 28 |
|  | R Supramarginal Gyrus, anterior division |  | 2.84 | 60 | -22 | 40 |
|  | Cluster 4 | 2,720 |  |  |  |  |
|  | R Middle Temporal Gyrus, anterior division |  | 5.41 | 56 | -2 | -20 |
|  | R Middle Temporal Gyrus, posterior division |  | 3.23 | 68 | -10 | -12 |
|  | Cluster 5 | 2,576 |  |  |  |  |
|  | L Inferior Temporal Gyrus, temporooccipital part |  | 3.43 | -50 | -58 | -20 |
|  | Cluster 6 | 2,512 |  |  |  |  |
|  | R Superior Frontal Gyrus |  | 4.80 | 24 | 30 | 48 |
|  | Cluster 7 | 1,208 |  |  |  |  |
|  | L Middle Temporal Gyrus, anterior division |  | 4.04 | -58 | -8 | -16 |
|  | L Middle Temporal Gyrus, posterior division |  | 4.29 | -64 | -12 | -14 |
|  | Cluster 8 | 1,016 |  |  |  |  |
|  | L Superior Frontal Gyrus |  | 3.51 | -26 | 30 | 52 |
|  | Cluster 9 | 984 |  |  |  |  |
|  | L Transverse Orbital Sulcus |  | 4.38 | -36 | 36 | -10 |
|  | Cluster 10 | 808 |  |  |  |  |
|  | L Planum Temporale |  | 4.21 | -52 | -38 | 14 |
|  | L Supramarginal Gyrus, posterior division |  | 2.93 | -54 | -48 | 18 |
|  | Cluster 11 | 808 |  |  |  |  |
|  | L Juxtapositional Lobule Cortex |  | 4.68 | -4 | -2 | 64 |
|  | Cluster 12 | 640 |  |  |  |  |
|  | R Middle Frontal Gyrus |  | 3.43 | 28 | 2 | 52 |
|  | Cluster 13 | 424 |  |  |  |  |
|  | L Superior Temporal Gyrus, posterior division |  | 3.29 | -64 | -28 | 2 |
|  | Cluster 14 | 312 |  |  |  |  |
|  | R Transverse Orbital Sulcus |  | 3.09 | 30 | 36 | -6 |
|  | Cluster 15 | 296 |  |  |  |  |
|  | R Supramarginal Gyrus, posterior division |  | 2.87 | 48 | -36 | 46 |
|  | Cluster 16 | 88 |  |  |  |  |
|  | R Inferior Temporal Gyrus, posterior division |  | 3.45 | 50 | -26 | -20 |
|  | Cluster 17 | 88 |  |  |  |  |
|  | R Subcallosal Cortex/Inferior Rostral Sulcus |  | 2.72 | 6 | 22 | -14 |
|  | Cluster 18 | 88 |  |  |  |  |
|  | L Supramarginal Gyrus, anterior division |  | 3.02 | -50 | -32 | 34 |
|  | Cluster 19 | 56 |  |  |  |  |
|  | L Inferior Temporal Gyrus, posterior division |  | 2.82 | -60 | -22 | -24 |
|  | Cluster 20 | 48 |  |  |  |  |
|  | L Middle Frontal Gyrus |  | 3.00 | -40 | 22 | 26 |
|  | Cluster 21 | 32 |  |  |  |  |
|  | R Planum Temporale |  | 3.00 | 40 | -34 | 12 |
|  | Cluster 22 | 24 |  |  |  |  |
|  | R Temporal Pole |  | 3.12 | 54 | 14 | -34 |
|  | Cluster 23 | 16 |  |  |  |  |
|  | R Planum Polare |  | 2.80 | 52 | -6 | -8 |
|  | Cluster 24 | 16 |  |  |  |  |
|  | L Insular Cortex |  | 2.65 | -32 | -26 | 4 |
|  | Cluster 25 | 16 |  |  |  |  |
|  | L Inferior Frontal Gyrus, pars triangularis |  | 2.74 | -38 | 32 | 12 |
|  | Cluster 26 | 8 |  |  |  |  |
|  | L Pallidum |  | 2.75 | -26 | -12 | -6 |
|  | Cluster 27 | 8 |  |  |  |  |
|  | L Occipital Pole |  | 3.03 | -12 | -90 | 12 |

^a^Within the Harvard-Oxford amygdala (*p*>.25), a total of 251 voxels (2,008 mm^3^) exceeded threshold. ^b^ 57% probability of lying within the Harvard-Oxford amygdala. ^c^ 42% probability of lying within the Harvard-Oxford amygdala. ^d^ 80% probability of lying within the Harvard-Oxford amygdala.

**Continued…**

**EXTENDED DATA FIG. 8-1**

**Detailed inferential statistics for the direct comparison of the BST to the Amygdala.**

| **Contrast**^a^ | **BST** | | **Amygdala** | | **R** | **Region × Condition** | **Cohen’s *d_z_*** | **TOST for a ‘Medium’ Effect**  ***d_Z_* = .35^a^** |
| --- | --- | --- | --- | --- | --- | --- | --- | --- |
|  | ***M*** | ***SD*** | ***M*** | ***SD*** |  |  |  |  |
| Uncertain Threat vs. Uncertain Safety | .2186 | .60067 | .1200 | .32312 | .273 | t(98)=1.637, p=.105 | .16 | t(98)=1.85, p=.034 |
| Certain Threat vs. Certain Safety | .2290 | .67035 | .2118 | .43576 | .300 | t(98)=.251, p=.802 | .03 | t(98)=3.23, p=.001 |
| Certain Threat vs. Uncertain Threat | -.2569 | .52219 | -.1997 | .35992 | .369 | t(98)=1.109, p=.270 | .11 | t(98)=2.37, p=.010 |

^a^ (Lakens, 2017).

**EXTENDED REFERENCE**

Lakens D (2017) Equivalence tests: A practical primer for t tests, correlations, and meta-analyses. Social Psychological and Personality Science 8:355-362.
